# Differential miRNA Expression Contributes to Emergence of Multiple Cancer Stem Cell Subpopulations in Human Colorectal Cancer

**DOI:** 10.1101/2023.02.06.527341

**Authors:** Victoria A. Stark, Caroline O. B. Facey, Lynn Opdenaker, Jeremy Z. Fields, Bruce M. Boman

## Abstract

One reason for lack of efficacy in cancer therapeutics is tumor heterogeneity. We hypothesize that tumor heterogeneity arises due to emergence of multiple cancer stem cell (CSC) subpopulations because miRNAs regulate expression of stem cell genes in CSCs. Our goal was to determine if: *i*) multiple CSC subpopulations exist in a human CRC cell population, and *ii*) miRNAs are differentially expressed in the different CSC subpopulations. We discovered that at least four different CSC populations (ALDH1, CD166, LGR5, LRIG1) exist in the HT29 cell line. CSC subpopulations were quantified using co-staining for multiple stem cell markers, isolated using FACS, and analyzed by NanoString miRNA profiling. The miRNA expression pattern in each CSC subpopulation was analyzed relative to miRNA expression patterns in other CSC subpopulations. Messenger RNAs predicted to be targeted by the upregulated miRNAs in each CSC subpopulation were: 1) identified using bioinformatics analyses, and 2) classified according to their predicted functions using David functional annotation analyses. We found multiple CSC subpopulations with a unique miRNA signature in each CSC subpopulation. Notably, the miRNAs expressed within one CSC subpopulation are predicted to target and downregulate the CSC genes and pathways that establish the other CSC subpopulations. Moreover, mRNAs predicted to be targeted by miRNAs in the different CSC subpopulations have different cellular functional classifications. That different CSC subpopulations express miRNAs that are predicted to target CSC genes expressed in other CSC subpopulations provides a mechanism that might explain the co-existence of multiple CSC subpopulations, tumor heterogeneity, and cancer therapy resistance.

## INTRODUCTION

The lack of effectiveness of many cancer treatments is attributed to tumor heterogeneity caused by co-existence of different tumor cell populations which are variably resistant to anti-cancer treatment [1,2]. We conjecture that different tumor cell populations are generated by different cancer stem cell (CSC) subpopulations, which gives rise to the tumor heterogeneity.

Consequently, we have been determining whether dysregulation of miRNAs might explain how the multiple CSC subpopulations emerge in CRC. For example, our previous data showed that: *i*) miRNA23b targets the CSC gene LGR5 [3]; *ii*) miRNA92a targets the CSC gene LRIG1 [4]; *iii*) CSC genes predicted to be targeted by miRNAs correlate with reduced levels of CSC gene expression [5]; *iv*) multiple subpopulations of CSCs exist in CRCs [6]. *Hypothesis*: tumor heterogeneity arises due to emergence of multiple CSC subpopulations because specific miRNAs target different SC genes in CSCs. If each different CSC subpopulation expresses miRNAs that target SC genes expressed in other CSC subpopulations, it would provide a mechanism that might explain the emergence of CSC subpopulations and tumor heterogeneity. Accordingly, in the current study, different CSC subpopulations in the HT29 CRC cell line were 1) identified using co-staining for multiple stem cell markers, 2) isolated using fluorescence activated cell sorting (FACS), 3) analyzed by NanoString miRNA profiling of miRNA expression patterns, and 4) evaluated using bioinformatics to classify cellular function.

## MATERIALS AND METHODS

### Cell Culture and Maintenance

HT29 cells were purchased from American Type Culture Collection (ATCC; Manassas, VA, USA) and grown to promote a single layer. Cells were cultured in McCoy’s 5A medium (GIBCO/Life Technologies) with 5% Fetal Bovine Serum (FBS) and 100 units/ml penicillin and 100 ug/ml streptomycin (1x Penicillin-Streptomycin). Cell cultures were maintained at 37 °C in humidified air at 5% CO_2_. Cells were grown in T-75 flasks (flow cytometric analysis) or T-175 flasks (Fluorescence Activated Cell Sorting) (VWR International; Bridgeport, NJ, USA) every 4-5 days until confluency. If cells were passaged every week with 1 mL suspension, cells were fed fresh media by the 3-4^th^ day. For experimentation, cells were cultured to about 50%. confluency.

### ALDEFLUOR Assay

#### ALDEFLUOR Assay

The ALDEFLUOR assay to identify and isolate ALDH+ cells was done as we previously described [3,7]. Briefly, the ALDEFLUOR kit was purchased from Stem Cell Technologies (Cambridge, MA) and preparation of reagents were carried out according to their protocol. Cells were grown for about 2-4 days until confluency of 50% was achieved. Every 48 hours the cell culture was given fresh media. The ALDEFLUOR kit contained the following: DEAB inhibitor, DMSO, HCl, and ALDEFLUOR Assay Buffer. The ALDEFLUOR kit and its reagents were left at room temperature at the start of the protocol. For ALDEFLUOR activation, DMSO (25 uL) was added to the ALDEFLUOR Reagent. Then the tube with DMSO was mixed at room temperature for 1 minute. Next, 25 uL of 2 N HCl was added and the tube was vortexed. The tube was incubated for 15 minutes at room temperature. Lastly, ALDEFLUOR Assay Buffer (360 uL) was added to the vial and mixed. The remaining ALDEFLUOR Reagent was stored in 20 uL aliquots at −20 °C for future experiments. Note: It is essential to keep the activated ALDEFLUOR reagent at 2-8 °C on ice during use.

#### ALDEFLUOR test for detection of ALDH+ cells

The assay was performed according to the Stem Cell Technologies published protocol with modification. Cells were grown to 50% confluency. Adherent cells were washed with PBS and detached from the flask using 0.25% Trypsin-EDTA (Fisher Scientific). After resuspension in fresh media, a cell count was performed to calculate the appropriate amount of suspension to give 1 million cells/mL. The appropriate amount from the calculation was transferred into 2 tubes labeled as control and sample. The cells were centrifuged at 500 x G for 5 minutes to pellet and then washed with 1 mL of PBS. The process of pipetting and pelleting was repeated once more. Then the PBS was aspirated, and cells were re-suspended in 1 mL of ALDEFLUOR assay buffer. For each sample, a control tube and sample tube were prepared. To the control tube, 5 uL of the DEAB inhibitor was added. To the sample tube, 5 uL of the activated ALDEFLUOR reagent was added and mixed with 1mL of the cell/ALDEFLUOR assay buffer suspension. Immediately after vortexing, 500 uL of the suspension was removed and added to the control tube with the inhibitor. Cells were then incubated for 25 minutes at 37 °C. After incubation, cells were pelleted by centrifugation, assay buffer aspirated, and cells re-suspended in 500 uL of fresh ALDEFLUOR buffer. Cells were then passed through a Falcon round bottom tube with a 50-μm cell strainer to obtain a single cell suspension (Corning, USA). Samples were placed on ice in the dark until ready for flow cytometric analysis via the BD LSR Fortessa.

### Flow Cytometry

HT29 cells were grown to 60% confluency and then lifted with Cell Stripper (Corning). After resuspension in fresh media, cells were calculated for 1 million cells per 1.5 mL tube. Cells were pelleted and then resuspended in 1 mL of Flow Cytometry Staining Buffer 1X (R&D Systems #FC001) for blocking. Cells were blocked for 1 hour on ice in the dark. After blocking, the cells were spun to aspirate the blocking buffer. The following primary antibodies were used: Anti-CD166 mouse monoclonal antibody conjugated with PE (phycoerythrin) at 5 uL (BD Biosciences #559263), anti-LRIG1 sheep polyclonal unconjugated antibody at 2.5 ug (Invitrogen #PA5-47928), and anti-LGR5 mouse monoclonal conjugated with PE at 1:100 (Origene #TA400001). Apart from the LGR5 antibody incubation (of 20 minutes), samples were incubated with antibodies for 30 minutes on ice in the dark.

The appropriate IgGs were also incorporated for the following primary antibodies. The following IgGs were included: mouse IgG1 – PE conjugate (5 uL for CD166; BD Biosciences #555749), purified sheep IgG (2 mg/mL for LRIG1 R&D Systems #5-001-A), and mouse IgG1-PE conjugate (10 uL for LGR5; R&D Systems #IC002P). It is worth noting that only the LRIG1 was incubated with a secondary antibody conjugated to allophycocyanin (APC; R&D Systems #F0127). LRIG1 cells were incubated with secondary antibody for 30 minutes in the dark on ice. Following antibody and IgG incubation, all cells were spun down to remove the antibody solution and resuspended in fresh flow cytometry staining buffer (R&D Systems #FC001). Cells were then passed through round bottom Falcon tubes for flow cytometric analysis using a BD LSR Fortessa.

### Fluorescence Activated Cell Sorting (FACS)

The following protocol for FACS utilized the flow cytometry protocol for isolating the different CSC populations. The amount of buffer solution and antibody concentrations were scaled up accordingly for successful sorting of ~7 million cells where two populations were analyzed. Once the protocol was completed, all cells were passed through the Falcon round bottom tube for FACS analysis via the BD FACS Aria II. After sorting, cells were spun down to remove supernatant. Next, 500 uL of Trizol was added to each tube and vortexed for 20 seconds. Lastly the tubes were set at room temperature in Trizol for 10 minutes. Following room temperature incubation, samples were transferred into 1.5 mL tubes, labeled, and stored at −80° C. For the combination sorts that involved the LGR5 antibody, sorted cells were centrifuged and placed in RNA Lysis Buffer from Zymo Research. The tubes were then immediately placed into the −80° C freezer for storage. Samples remained in storage until each set with appropriate replicates were complete and ready to be sent for NanoString miRNA profiling at the Wistar Genomics Facility (Philadelphia, PA).

### miRNA Profiling

The data received from the Genomics Core Facility at the Wistar Institute was then analyzed using the nSolver 4.0 from NanoString Technologies (Seattle, WA). Heat maps were generated for the TOP 50 miRNAs expressed for each of the different sort combinations of LRIG1, LGR5, CD166, and ALDH. For clarification, the sort combinations were done in pairs of two during FACS and NanoString profiling analysis. We then generated a list of ratios for each pair of stem cell subpopulations and sorted the list according to their respective *p*-values. Only statistically significant (*p*<0.05) differentially expressed miRNAs were selected. The first 50 miRNAs with the highest expression levels were selected and used to generate the heatmaps presented in this paper.

### Statistical Analysis

T-tests and its corresponding p-values for the different miRNAs were generated by the nSolver 4.0 software. The interpretation of the different miRNA p-values helped designate our confidence in the miRNAs that are said to target LRIG1, CD166, LGR5, and ALDH, respectively.

### Bioinformatics Analysis

Bioinformatics analysis was done using the TCGA database [Pan, C-Y and Lin, W-C; miR-TV Introduction available online] and http://www.miRbase.org to look for individual miRNAs and the relevant clusters that are targeted in stem cell genes. In addition to the targeted SC genes, we analyzed the miRNA expression level and different clinical correlations with CRC. Lastly, we also investigated miRNA isoform (isomiR) expression as it displayed variability in expression regarding the cancer type. Additional databases used were: i) miRBase for identifying miRNA clusters [miRbase Available online], TargetScan for identifying isomiRs and predicted miRNA targets [TargetScanHuman 7.2 Available online], and GeneCards for identifying miRTarBase predicted miRNA targets [GeneCards – Human Genes | Gene Database | Gene Search Available online]. The DAVID functional annotation tool was used to classify the different mRNA functions.

## RESULTS

Our approach was to use flow cytometric analysis to determine the proportion of different CSC subpopulations in the HT29 CRC cell line. The bar graph shown in Figure 1 gives the percentage of CSCs in the HT29 CRC cell population that stain for the different SC markers: ALDH+ (59.3%), LRIG1+ (40.2%), CD166+ (25.2%), and LGR5+ (10.2%) cells. This analysis raised the question: Can specific CSC subpopulations be identified by analyses for the expression of multiple different SC markers?

**Figure 1.**
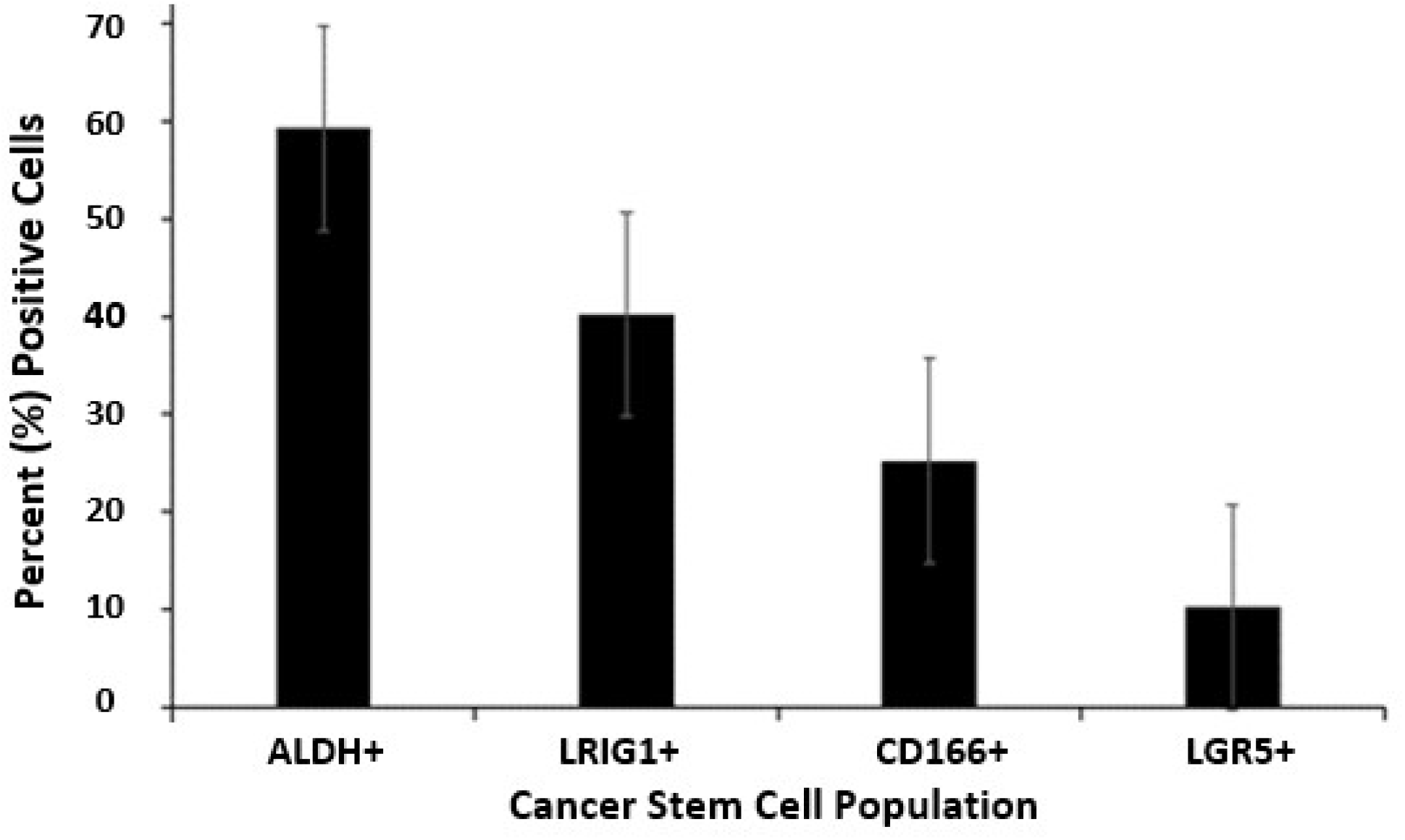
Proportion of different CSC subpopulations in the HT29 CRC cell line. The bar graph gives the percent ALDH+, LRIG1+, CD166+, and LGR5+ cells in the HT29 CRC cell population as determined by flow cytometric analysis. Error bars represent standard error of the mean (n = 3-7 experimental replicates per marker).

We then evaluated six different combinations of SC marker pairs: CD166 & ALDH, LRIG1 & CD166, LRIG1 & ALDH, LGR5 & LRIG1, LGR5 & CD166, LGR5 & ALDH (described in Materials & Methods). Thus, in each of the different experiments, co-staining for two different SC markers was analyzed which led to the isolation of four different cell subpopulations: cells that are co-positive for both markers (double positive), cells that are positive for just one marker or the other marker (two different populations positive for a single marker), and cells that are negative for both markers (double negative). The miRNA expression in each cell subpopulation was then analyzed using NanoString profiling to identify miRNAs that are differentially expressed in the different CSC subpopulations (Table S1). The genes (mRNAs) that were predicted to be targeted by differentially expressed miRNAs in each CSC subpopulation were then identified, and the function of these mRNAs was classified using bioinformatics analysis (see Methods).

Results from the LGR5 and ALDH sort are presented in Figure 2. The results on the remaining CSC subpopulations are in the supplementary data section (Supplemental Figures S1-S5). The different functional classifications of the mRNAs predicted to be targeted by the miRNAs in each cell subpopulation are given in Table S2. We also did a more detailed analysis of the results from the LGR5 & ALDH sort because it had the lowest size (<1%) of the co-positive (LGR5+/ALDH+) cell subpopulation, and one of the highest numbers of differentially expressed miRNAs (Table S1). In view of underlying mechanisms, these findings might reflect to differences in functionality between LGR5+ cells and ALDH+ cells and the function of the genes that are targeted by miRNAs in the different subpopulations.

**Figure 2.**
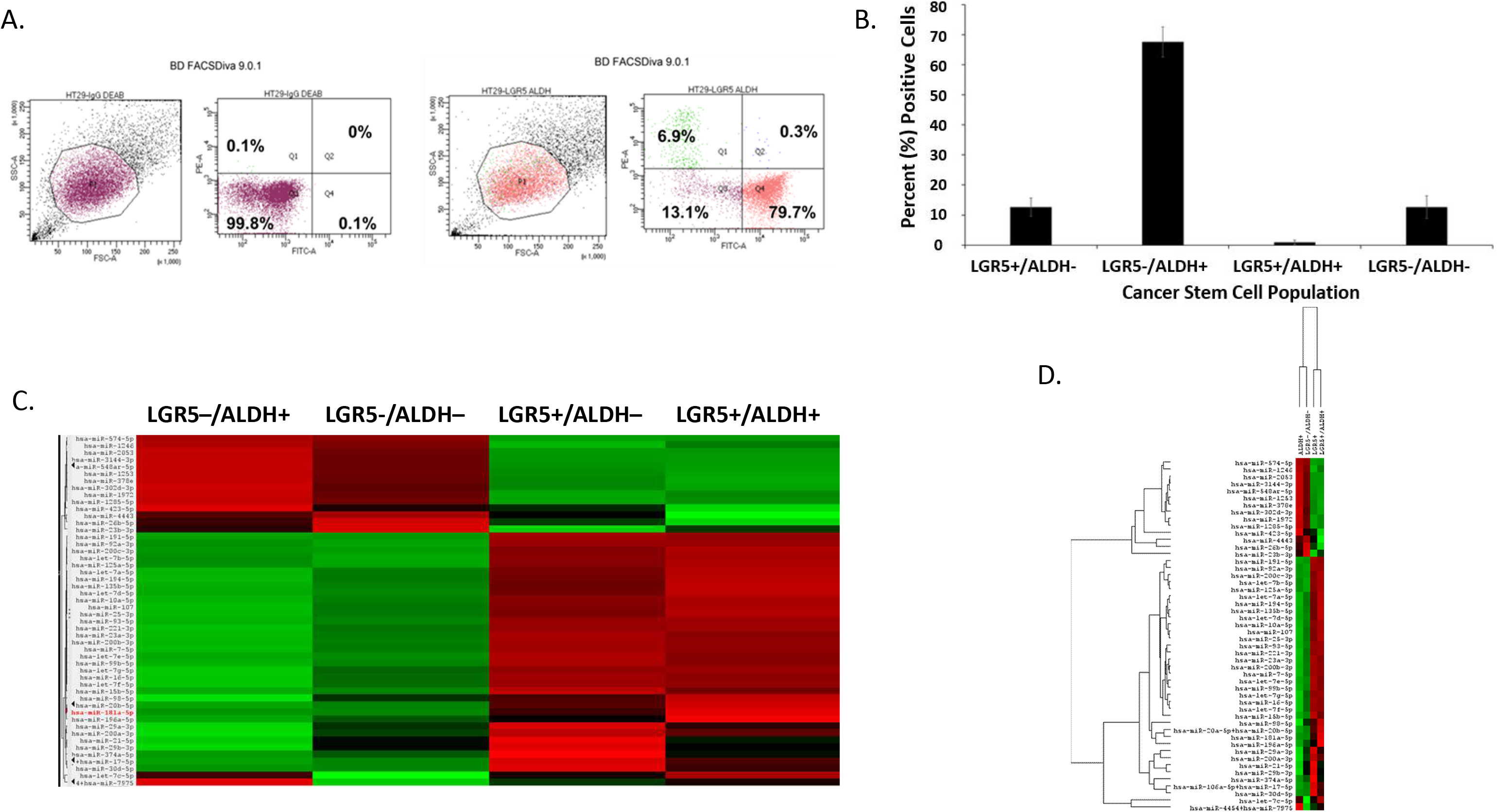
Proportion of different CSC subpopulations from FACS analysis of LGR5 and ALDH marker expression. This figure gives results from FACS analysis of stained CRC cells using a combination of the SC marker pair ALDH & LGR5. *<u>Panel A</u>* gives a representative dot blot graph illustrating the proportion of cells in LGR5+/ALDH–, ALDH+/LGR5–, LGR5+/ALDH+, and LGR5–/ALDH– cell subpopulations. *<u>Panel B</u>* gives a bar graph showing the average percentage positive cells in each of the four different CSC subpopulations. Error bars represent standard error of the mean. *<u>Panel C</u>* gives a heatmap from NanoString profiling analysis showing differential expression of the top 50 miRNAs in the four different isolated CSC subpopulations (increased expression = green; decreased expression = red for each CSC marker). NanoString miRNA profiling. *<u>Panel D</u>* provides the same heatmap with the respective dendrogram. Panels C and D visually illustrate the patterns that are seen when a large set of miRNAs is surveyed and they are not meant to show details, which is why the vertical axes are not legible.

The proportions of the different cell subpopulations (LGR5+/ALDH–, ALDH+/LGR5–, LGR5+/ALDH+, LGR5–/ALDH–) are shown as a FACS dot plot and bar graph in Figures 2A & 2B. Differential expression of the miRNAs is shown as a heatmap in Figures 2C & 2D. The top 10 miRNAs as ranked by *p*-value in the LGR5+/ALDH– subpopulation and in the ALDH+/LGR5– subpopulation are listed in Figures 3 & 4, respectively. The mRNAs predicted to be targeted by these miRNAs were identified using bioinformatics analysis and the functional classification of these mRNAs is provided in the pie chart graphs in Figures 3 & 4. The functional classification of mRNAs predicted to be targeted by miRNAs identified in LGR5+/ALDH– cells was transcriptional regulation and zinc finger motifs (Figure 3). In contrast, the classification of predicted mRNAs in ALDH+/LGR5– cells was phosphoproteins and protein binding (Figure 4).

**Figure 3.**
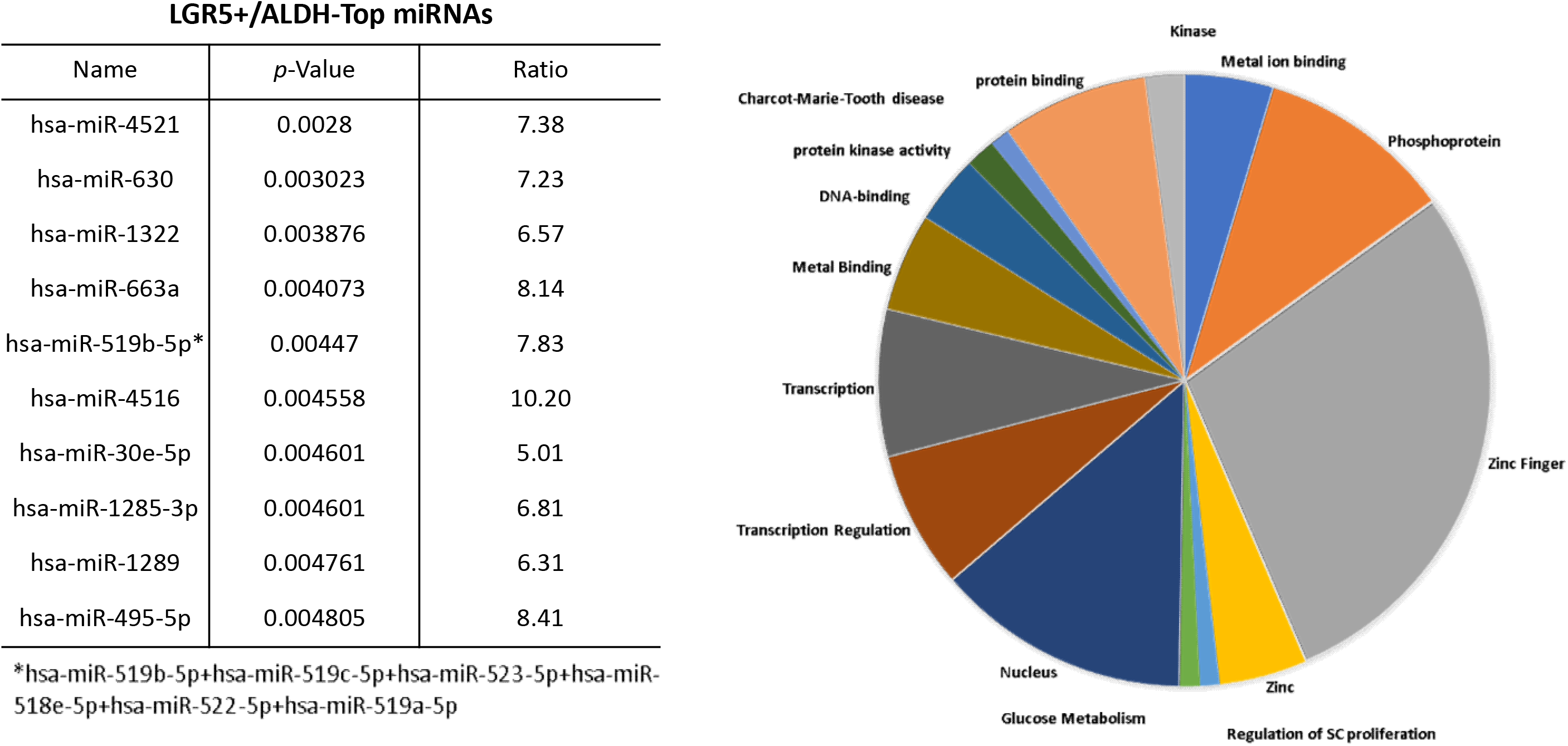
The top 10 miRNAs and functional analysis of predicted targeted genes in LGR5+/ALDH– cells. The table lists the top 10 miRNAs expressed in LGR5+/ALDH– cells. miRNAs were ranked according to *p*-value <0.05 for statistical significance. The asterisk depicts the original name for the miRNA that was abbreviated in the Table. The pie chart shows the functional classification identified by David analysis of the mRNAs predicted to be targeted by the miRNAs expressed in LGR5+/ALDH– cells. A summary of the functions of mRNAs predicted to be targeted by the miRNAs is given in Table S2.

**Figure 4.**
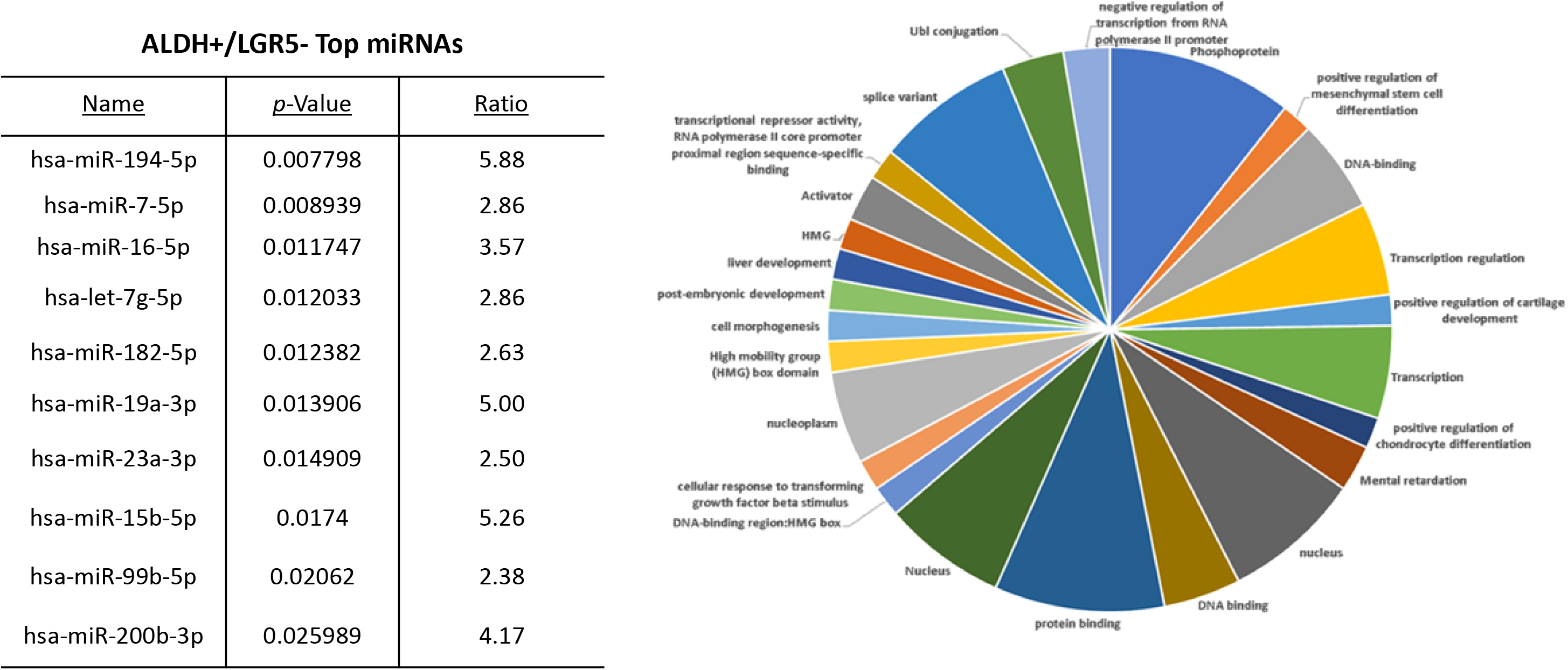
The top 10 miRNAs and functional analysis of predicted targeted genes in ALDH+/LGR5– cells. The table lists the top 10 miRNAs expressed in ALDH+/LGR5– cells. miRNAs were ranked according to *p*-value <0.05 for statistical significance. The pie chart shows the functional classification identified by David analysis of the mRNAs predicted to be targeted by the miRNAs expressed in ALDH+/LGR5– cells. A summary of the functions of mRNAs predicted to be targeted by the miRNAs is given in Table S2.

We further analyzed results from the LGR5 & ALDH sort to determine what might distinguish the different CSC subpopulations from one another. We did this using two different approaches. First, we analyzed the top 10 miRNAs ranked based on level of upregulation of miRNA expression in the LGR5+/ALDH– subpopulation versus ALDH+/LGR5– subpopulation. Bioinformatics analysis was then used to determine which genes in the RA and WNT signaling pathways (Tables S2 & S3) were predicted to be targeted by these miRNAs. We found that 35 mRNAs in the RA signaling pathway are predicted to be targeted by the top miRNAs expressed in LGR5+/ALDH– cells (Figure 4) and 18 mRNAs in the WNT signaling pathway are predicted to be targeted by the top miRNAs expressed in ALDH+/LGR5– cells (Figure 5).

**Figure 5.**
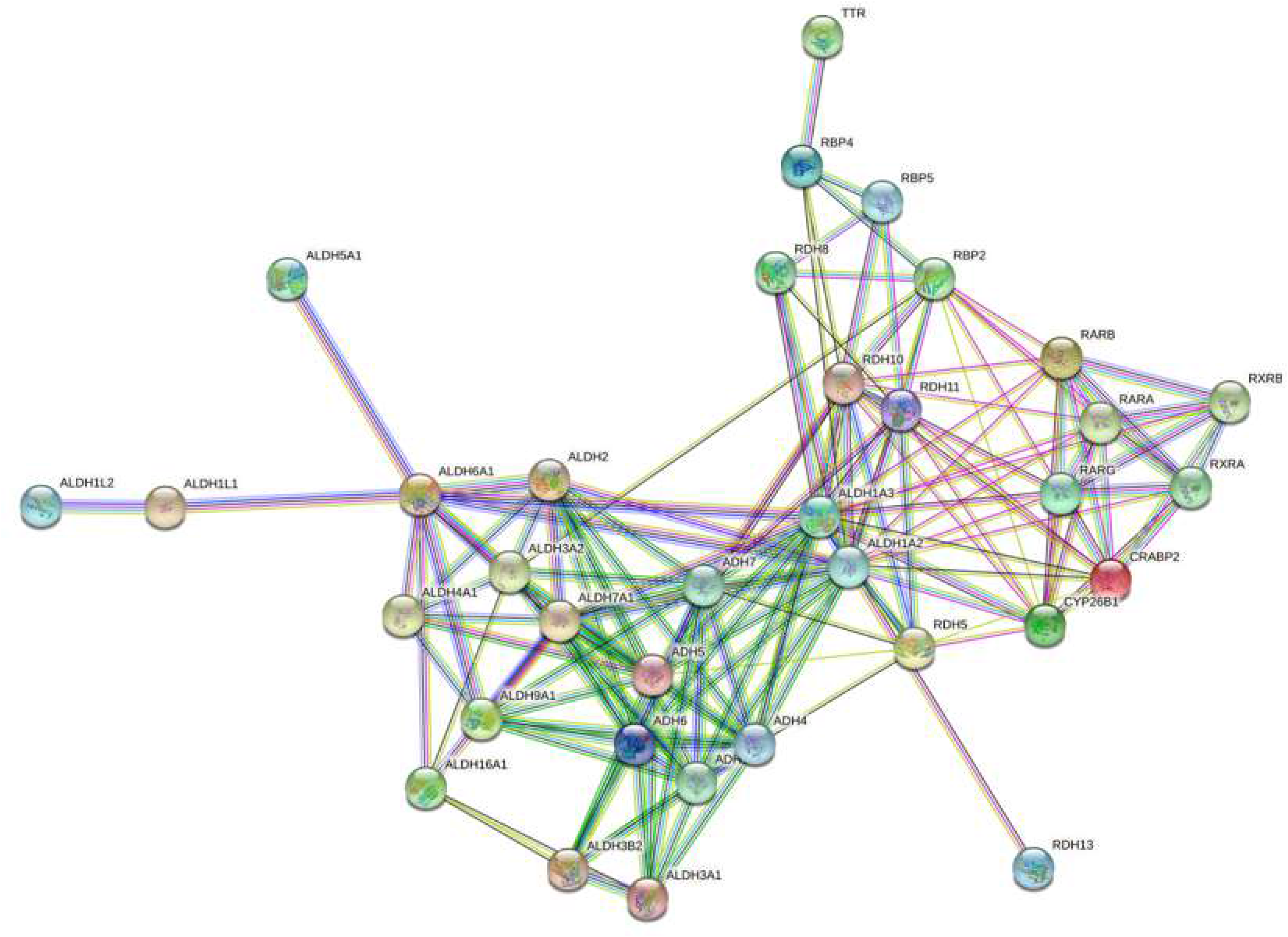
A string network map of genes in the retinoic acid signaling pathway that are predicted to be targeted by miRNAs expressed in LGR5+/ALDH– cells. The following 25 mRNAs are predicted to be targeted by miRNAs that are selectively expressed in LGR5+ cells (not expressed in ALDH+ cells): *ADH1B, ADH4, ADH5, ADH6, ADH7, ALDH16A1, ALDH1A3, ALDH1L1, ALDH2, ALDH3A1, ALDH3A2, ALDH3B2, ALDH4A1, ALDH5A1, ALDH7A1, ALDH9A1, CRABP2, RBP2, RBP4, RBP5, RDH11, RDH13, RDH5, RDH8, RXRB*.

**Figure 6.**
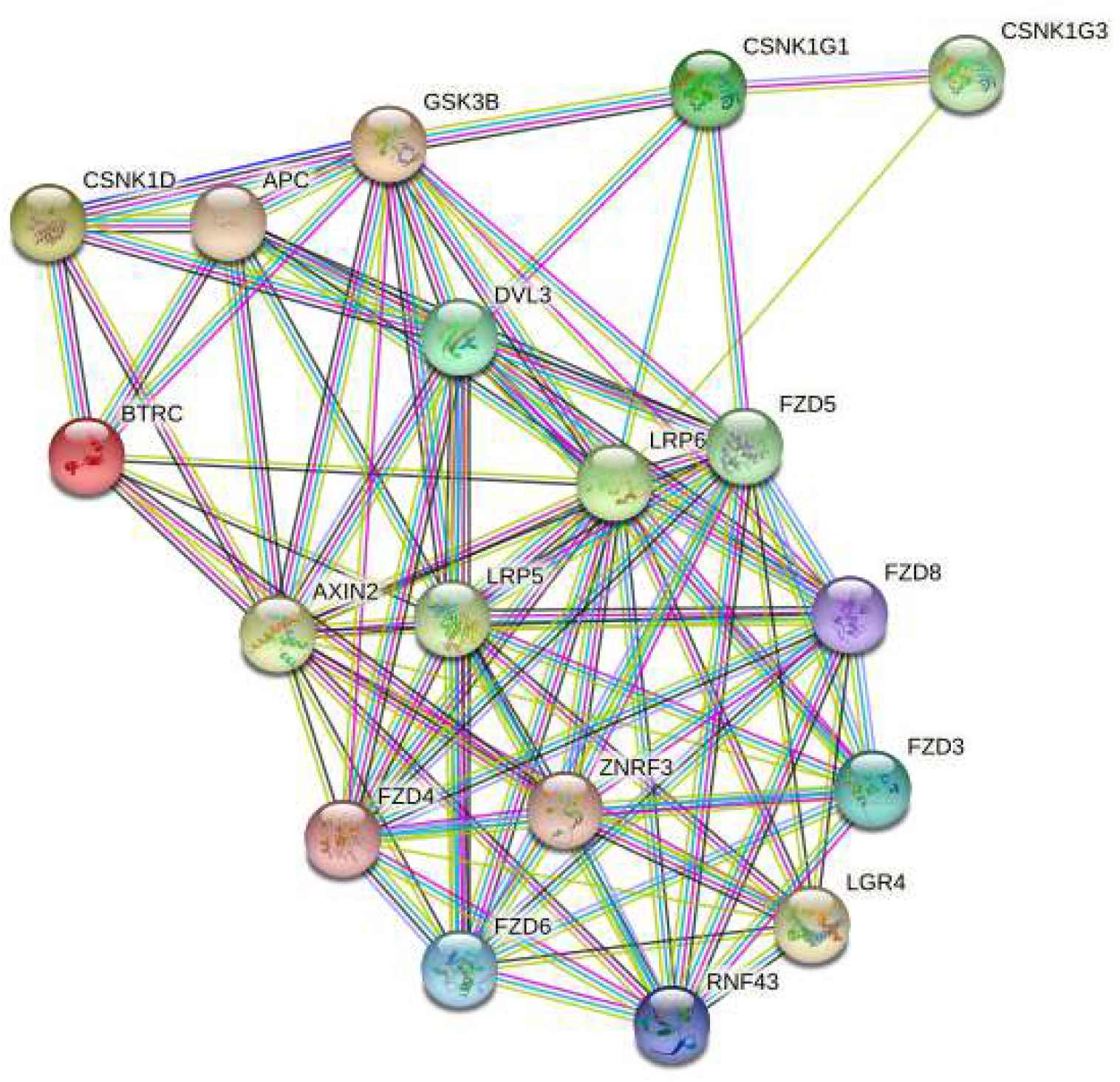
A string network map of genes in the WNT signaling pathway that are predicted to be targeted by miRNAs expressed in ALDH+/LGR5– cells. Two mRNAs in the WNT signaling pathway (*AXIN2* & *LRP5*) are predicted to be targeted by miRNAs selectively expressed in ALDH+ cells (not expressed in LGR5+ cells).

In our second approach, we analyzed miRNAs that are selectively expressed in each CSC subpopulation. Specifically, we analyzed miRNAs that are expressed in LGR5+/ALDH– cells but not expressed in ALDH+/LGR5– cells, and vice versa. This analysis showed that in the RA signaling pathway, 25 mRNAs (mostly dehydrogenases) are predicted to be targeted by miRNAs selectively expressed in LGR5+/ALDH– cells. We also found that 2 mRNAs in the WNT signaling pathway are predicted to be targeted by miRNAs selectively expressed in ALDH+/LGR5– cells. The miRNAs that were predicted to selectively target LRP5/6, AXIN2, and the dehydrogenases from ALDH & LGR5 experiment were further analyzed to see if these miRNAs are expressed in CSC subpopulations isolated from our other FACS sorting experiments (Table 2). Indeed, we identified an miRNA signature for ALDH+ cells (miR-16-5p, miR-23a-3p, miR15b-5p, miR-15a-5p, miR375, miR107), and for LGR5+ cells (miR-4521, miR-630, miR-1322, miR-519b-5p, miR-4516, miR-1285-3p, miR-1289, miR-495-5p). Thus, based on miRNA expression in CSC subpopulations from our other FACS sorts, we found that there are specific miRNA signatures that characterize ALDH+ cells and LGR5+ cells.

## DISCUSSION

The key findings in our study are: 1) miRNAs are selectively expressed in different CSC subpopulations; 2) miRNAs that are predicted to target and downregulate genes that encode CSC proteins in one CSC subpopulation are SC genes likely upregulated in the other CSC subpopulation. These findings support our hypothesis that tumor heterogeneity arises due to emergence of multiple CSC subpopulations because specific miRNAs target different SC genes in CSCs.

Using flow cytometry and FACS analyses, we found that four different CSC subpopulations were present in each experiment: two single CSC populations, a co-positive and a co-negative population. The presence of 4 subpopulations shows that intra-tumoral CSC heterogeneity exists in a CRC cell line. We found that, among the different experiments, the LGR5 & ALDH sort had the lowest co-positive staining, indicating that LGR5 and ALDH mark distinctly different CSC subpopulations.

We next determined, using NanoString profiling, which miRNAs are differentially expressed between the different CSC subpopulations, and identified mRNAs predicted to be targeted by the upregulated miRNAs using additional bioinformatics tools. Notably, a unique miRNA signature was identified for each CSC subpopulation. The mRNAs predicted to be targeted by the miRNAs in each CSC subpopulation were found to have very different functional classifications (Table S2). Notably, bioinformatics analyses of NanoString profiling results showed that many targeted mRNAs play important functional roles in maintaining properties of stemness. For example, the miRNAs that are up in the ALDH+/LGR5– subpopulation are predicted to target and suppress expression of genes in WNT signaling. And, these WNT pathway genes are predicted to be upregulated in the LGR5+/ALDH– subpopulation.

Conversely, the miRNAs that are up in LGR5+/ALDH– subpopulation are predicted to target and downregulate expression of genes (dehydrogenases) involved in retinoic acid (RA) signaling. And, these RA pathway genes are predicted to be upregulated in the ALDH+/LGR5– subpopulation. Thus, our results indicate that the miRNAs expressed in LGR5+/ALDH– CSCs decrease the expression of mRNAs that encode proteins essential for the existence of ALDH+/LGR5– CSCs, and vice versa.

Components of RA signaling were evaluated because RA signaling appears to mainly occur through ALDH+ SCs [7,8]. Indeed, ALDH is a key component in RA pathway and ALDH is a key SC marker that can track CSC overpopulation during CRC development [9]. Components of WNT signaling were also considered because over 90% of CRC patients have mutations in the WNT pathway [10]. It is constitutively activated WNT signaling, due to *APC* mutations, that is the main driver of CRC growth and development [11,12]. The WNT signaling pathway is also important as LGR5 is a receptor for R-spondins and is a key factor in the canonical WNT signaling pathway [13]. Notably, the WNT pathway incorporates signaling via LGR5 in CRC growth. Accordingly, we evaluated the different mRNAs in the WNT and RA signaling pathways that are predicted to be targeted by top miRNAs in LGR5+/ALDH– cells and in ALDH+/LGR5– cells.

Thus, investigating the role of miRNAs in regulation of RA and WNT signaling is a logical step to further understand how dysregulated miRNAs contribute to emergence of different CSC subpopulations. Accordingly, we took a bioinformatics approach to investigate predicted functions for the mRNAs targeted by the miRNAs in LGR5+/ALDH– and ALDH+/LGR5– cells. We surmised that RA and WNT signaling pathways are likely downregulated by miRNAs in LGR5+/ALDH– and ALDH+/LGR5– cells, respectively.

In our bioinformatics analysis of miRNAs in LGR5+/ALDH– cells, we discovered that many of these miRNAs are predicted to target expression of a number of dehydrogenases in the RA signaling pathway. These dehydrogenases included aldehyde dehydrogenase (ALDH), alcohol dehydrogenase (ADH), and retinol dehydrogenase (RDH). While LGR5 plays a key role in WNT signaling, its role in RA signaling is largely unknown. However, our previous studies indicated that the WNT and RA signaling pathways are linked and play a role in CRC development [14]. Thus, it is possible that RA signaling and WNT signaling are inversely correlated to each other, and have opposing functions that maintain co-existence of colonic SC populations and homeostasis of colon tissues. Other studies show RA receptors can induce or downregulate WNT/β-catenin signaling during chondrocyte development [15] and development of cardiac/skeletal muscle [16]. RA signaling and WNT signaling also have opposing functional roles that control the development of different parts of the embryo [17,18].

In our bioinformatics analysis of miRNAs in ALDH+/LGR5– cells, we discovered that many of these miRNAs are predicted to target expression of AXIN2 and LRP5, which are components of the WNT signaling pathway. The WNT pathway mRNAs targets were of particular interest to evaluate because LGR5 is known to promote WNT/β-catenin signaling in the SC origin of CRCs [19]. In CRC, LRP5/6 plays an important role in WNT signaling as the WNT ligands bind to LRP5/6 and Frizzled receptors to promote WNT signaling via other WNT pathway components such as AXIN2 and β-catenin [20]. Indeed, AXIN2 is classified as a tumor suppressor gene, and AXIN2 germline mutations occur in hereditary CRC patients and predispose them to develop CRC [21]. Moreover, as noted above, LRP5 acts as a receptor for WNT ligands [20]. Thus, our findings indicate that cross regulatory miRNA-based mechanisms control expression of WNT and RA signaling components that leads to emergence of different CSC subpopulations in CRC tissues.

## CONCLUSIONS

Our results provide key information on a mechanism that explains how miRNA expression plays a role in the emergence of multiple CSC subpopulations that contributes to tumor heterogeneity during CRC development. Specifically, our findings indicate that multiple CSC subpopulations emerge in HT29 CRC cells due to the expression of unique miRNAs that target different SC genes (and their co-expressed genes), resulting in intra-tumoral CSC heterogeneity. Thus, our research study has vast clinical significance because it provides insight into how dysregulation of miRNA expression leads to emergence of CSC subpopulations in CRC, which advances our understanding of what causes tumor heterogeneity. Our findings also provide clues as to how to use miRNA-based drugs to specifically target and eliminate CSCs and improve efficacy of anti-cancer therapies [22–25].

**Table 1.** MicroRNA expression patterns in ALDH+ cells and LGR5+ cells in CSC sub-populations isolated from different FACS sorting experiments.

|  |  |  |  |  |
| --- | --- | --- | --- | --- |
|  | **LGR5/ALDH Sort** | **LGR5/CD166 Sort** | **LGR5/LRIG1 Sort** | **ALDH/LRIG1 Sort** |
|  | **ALDH+ Cells** | **LGR5+ Cells** | **LGR5+ Cells** | **ALDH+ Cells** |
| miR-16-5p | ↑ | ↓ | ↓ | ↑ |
| miR-23a-3p | ↑ | ↓ | ↓ | ↑ |
| miR-15b-5p | ↑ | ↓ | ↓ | ↑ |
| miR-15a-5p | ↑ | ↓ | ↓ | ↑ |
| miR-375 | ↑ | ↓ | ↑ | ↑ |
| miR-107 | ↑ | ↑ | ↓ | ↑ |
|  | **LGR5+ Cells** | **LGR5+ Cells** | **LGR5+ Cells** | **ALDH+ Cells** |
| miR-4521 | ↑ | ↑ | ↑ | ↓ |
| miR-630 | ↑ | ↑ | ↑ | ↓ |
| miR-1322 | ↑ | ↑ | ↑ | ↓ |
| miR-519b-5p | ↑ | ↑ | ↑ | ↓ |
| miR-4516 | ↑ | ↑ | ↑ | ↓ |
| miR-1285-3p | ↑ | ↑ | ↑ | ↓ |
| miR-1289 | ↑ | ↑ | ↑ | — |
| miR-495-5p | ↑ | ↑ | ↑ | ↓ |

## Acknowledgements

A special thanks to Dr. Erica M. Selva for her leadership of the graduate program as well as for her dedication and support of this research project. We thank Molly Lausten, Brian Osmond and Chi Zhang for helpful comments and valuable suggestions. The authors also thank Dr. Nicholas Petrelli for his support of the project that was conducted at the Center for Translational Cancer Research and we thank the Genomics Facility at the Wistar Institute for Nanostring profiling.

## SUPPLEMENTAL FIGURE LEGENDS

**Figures S1-S5** give results from FACS isolation and NanoString profiling analysis of stained CRC cells using five different combinations of SC marker pairs: <u>S1</u>) CD166 & ALDH, <u>S2</u>) LRIG1 & CD166, <u>S3</u>) LRIG1 & ALDH, <u>S4</u>) LGR5 & LRIG1, <u>S5</u>) LGR5 & CD166. In each figure, *<u>Panel A</u>* gives a FACS dot blot graph from a representative experiment showing proportion of the cells in the four different cell subpopulations: cells that are co-positive for both markers (double positive), cells that are positive for just one marker or the other marker (two different populations positive for a single marker), and cells that are negative for both markers (double negative). *<u>Panel B</u>* gives a bar graph that shows the average percentage positive cells in each of these four different CSC subpopulations. Error bars represent standard error of the mean. *<u>Panel C</u>* gives a heatmap showing differential expression of the top 50 miRNAs in the four different isolated CSC subpopulations (increased expression = green; decreased expression = red for each CSC marker). *<u>Panel D</u>* shows the same heatmap with the corresponding dendrogram. Panels C and D visually illustrate the patterns that are seen when a large set of miRNAs is surveyed and they are not meant to show details, which is why the vertical axes are not legible. *<u>Panel E</u>* gives a table listing the top 10 miRNAs expressed in each subpopulation. *<u>Panel F</u>* gives a pie chart graph showing the functional classification identified by David analysis of the mRNAs predicted to be targeted by the miRNAs expressed in each subpopulation. In each analysis, the top 10 miRNAs were ranked by *p*-value (*p* < 0.05) in order to determine the mRNAs that are predicted to be targeted by miRNAs in each subpopulation. Table S2 gives a summary of the functions of mRNAs predicted to be targeted by the miRNAs.

## Notes

**Financial support**: This study was supported in part by The Lisa Dean Moseley Foundation (BB), Cancer B*Ware Foundation (BB), The Cawley Center for Translational Cancer Research Fund (VS, CF, BB), University of Delaware Department of Biological Sciences (VS), and INBRE NIH/NIGMS GM103446 (LO).

### Competing Interest Statement

The authors have declared no competing interest.

