## Supplemental Tables 1-4 and Figures 1-5 for "Differential miRNA Expression Contributes to Emergence of Multiple Cancer Stem Cell Subpopulations in Human Colorectal Cancer"

### SUPPLEMENTAL INFORMATION

**Table S1. Significantly expressed miRNAs in different CSC subpopulations*.**

| LGR5/ALDH FACS Isolation | miRNAs |
|---|---|
| LGR5+/ALDH– cells | 310 |
| LGR5–/ALDH+ cells | 15 |
| CD166/ALDH FACS Isolation | miRNAs |
| CD166+/ALDH– cells | 1 |
| CD166–/ALDH+ cells | 2 |
| CD166/LRG5 FACS Isolation | miRNAs |
| CD166+/LRG5– cells | 1 |
| CD166–/LRG5+ cells | 102 |
| LRIG1/LGR5 FACS Isolation | miRNAs |
| LRIG1+/LGR5– cells | 4 |
| LRIG1–/LGR5+ cells | 673 |
| LRIG1/ALDH FACS Isolation | miRNAs |
| LRIG1+/ALDH– cells | 101 |
| LRIG1–/ALDH+ cells | 4 |
| CD166/LRIG1 FACS Isolation | miRNAs |
| CD166+/LRIG1– cells | 29 |
| CD166–/LRIG1+ cells | 4 |

*Based on $p$-value < 0.05.

| **Table S2. The mRNAs predicted to be targeted by the miRNAs in each CSC subpopulation were found to have different functional classifications** | | | | |
|---|---|---|---|---|
| LGR5 vs. ALDH | zinc finger motifs | *vs.* | phosphoproteins, and protein binding | Figure 2 vs. Figure 3 |
| CD166 *vs*. ALDH | alternative splicing, phosphoproteins, and membrane | *vs.* | phosphoprotein, nucleus, cytoplasm, and nucleus functions | Figure S2 *vs*. Figure S3 |
| LRIG1 *vs*. CD166 | alternative splicing, transcriptional regulation, and nucleus | *vs.* | phosphoprotein functions | Figure S5 *vs*. Figure S6 |
| LRIG1 *vs*. ALDH | alternative splicing and as protein kinase domains | *vs.* | phosphoproteins, protein binding | Figure S8 *vs*. Figure S9 |
| LGR5 *vs*. LRIG1 | cytoplasm and different protein domains | *vs.* | alternative splicing, protein binding, and phosphoprotein | Figure S11 *vs*. Figure S12 |
| LGR5 *vs*. CD166 | splice variant, phosphoprotein and zinc finger motifs | *vs.* | phosphoprotein, protein binding, cytoplasm, and nucleus | Figure S14 *vs*. Figure S15 |

**Tables S3 & S4. BIOINFORMATICS ANALYSIS OF LGR5 and ALDH DATA**

To determine what might distinguish the different CSC subpopulations from each other, a detailed analysis of the LGR5 and ALDH data was done. Then a bioinformatics analysis was done to determine which genes in the Retinoic Acid (Table 1) and WNT (Table 2) signaling pathways were predicted to be targeted by these miRNAs.

| **Table S3. Retinoic Acid Pathway Genes Analyzed** | |
|---|---|
| *ADH1A* | Alcohol Dehydrogenase 1A |
| *ADH1B* | Alcohol Dehydrogenase 1B |
| *ADH1C* | Alcohol Dehydrogenase 1C |
| *ADH4* | Alcohol Dehydrogenase 4 |
| *ADH5* | Alcohol Dehydrogenase 5 |
| *ADH6* | Alcohol Dehydrogenase 6 |
| *ADH7* | Alcohol Dehydrogenase 7 |
| *ALDH16A1* | Aldehyde Dehydrogenase 16A1 |
| *ALDH18A1* | Aldehyde Dehydrogenase 18A1 |
| *ALDH1A1* | Aldehyde Dehydrogenase 1A1 |
| *ALDH1A2* | Aldehyde Dehydrogenase 1A2 |
| *ALDH1A3* | Aldehyde Dehydrogenase 1A3 |
| *ALDH1B1* | Aldehyde Dehydrogenase 1B1 |
| *ALDH1L1* | Aldehyde Dehydrogenase 1L1 |
| *ALDH1L2* | Aldehyde Dehydrogenase 1L2 |
| *ALDH2* | Alcohol Dehydrogenase 2 |
| *ALDH3A1* | Aldehyde Dehydrogenase 3A1 |
| *ALDH3A2* | Aldehyde Dehydrogenase 3A2 |
| *ALDH3B1* | Aldehyde Dehydrogenase 3B1 |
| *ALDH3B2* | Aldehyde Dehydrogenase 3B2 |
| *ALDH4A1* | Aldehyde Dehydrogenase 4A1 |
| *ALDH5A1* | Aldehyde Dehydrogenase 5A1 |
| *ALDH6A1* | Aldehyde Dehydrogenase 6A1 |
| *ALDH7A1* | Aldehyde Dehydrogenase 7A1 |
| *ALDH8A1* | Aldehyde Dehydrogenase 8A1 |
| *ALDH9A1* | Aldehyde Dehydrogenase 9A1 |
| *CRABP1* | Cellular Retinoic Acid Binding Protein 1 |
| *CRABP2* | Cellular Retinoic Acid Binding Protein 2 |
| *CRBP* | Cellular Retinol Binding Protein |
| *CYP26A1* | Cytochrome P450 26A1 |
| *CYP26B1* | Cytochrome P450 26B1 |
| *CYP26C1* | Cytochrome P450 26C1 |
| *RARA* | Retinoic Acid Receptor Alpha |
| *RARB* | Retinoic Acid Receptor Beta |

| | |
|---|---|
| *RARG* | Retinoic Acid Receptor Gamma |
| *RBP1* | Retinol Binding Protein 1 |
| *RBP2* | Retinol Binding Protein 2 |
| *RBP3* | Retinol Binding Protein 3 |
| *RBP4* | Retinol Binding Protein 4 |
| *RBP5* | Retinol Binding Protein 5 |
| *RBP7* | Retinol Binding Protein 7 |
| *RDH10* | Retinol Dehydrogenase 10 |
| *RDH11* | Retinol Dehydrogenase 11 |
| *RDH12* | Retinol Dehydrogenase 12 |
| *RDH13* | Retinol Dehydrogenase 13 |
| *RDH14* | Retinol Dehydrogenase 14 |
| *RDH16* | Retinol Dehydrogenase 16 |
| *RDH5* | Retinol Dehydrogenase 5 |
| *RDH8* | Retinol Dehydrogenase 8 |
| *RXRA* | Retinoid X Receptor Alpha |
| *RXRB* | Retinoid X Receptor Beta |
| *RXRG* | STRA6 |
| *STRA6* | Stimulated by retinoic acid 6 |
| *TTR* | Transthyretin |

| **Table S4. WNT Pathway Genes Analyzed** | |
|---|---|
| *APC* | adenomatous polyposis coli |
| *AXIN1* | aka Axin 1 a protein coding gene |
| *AXIN2* | aka Axin 2 a protein coding gene |
| *BTRC* | Beta-transducin repeat containing E3 ubiquitin protein ligase |
| *CSNK1A1/CK1* | casein kinase 1 alpha 1 |
| *CK1 beta* | casein kinase 1 beta 1 |
| *CK1 gamma 1* | casein kinase 1 gamma 1 |
| *CK1 gamma 2* | casein kinase 1 gamma 2 |
| *CK1 gamma 3* | casein kinase 1 gamma 3 |
| *CK1 delta* | casein kinase 1 delta |
| *CK1 epsilon* | casein kinase 1 epsilon |
| *CTNNB1* | Beta-catenin 1 |
| *DVL1* | dishevelled (DSH) DVL in mammals 1 |
| *DVL2* | dishevelled (DSH) DVL in mammals 2 |
| *DVL3* | dishevelled (DSH) DVL in mammals 3 |
| *FZD1* | Frizzled Class Receptor 1 |
| *FZD2* | Frizzled Class Receptor 2 |
| *FZD3* | Frizzled Class Receptor 3 |

| | |
|---|---|
| *FZD4* | Frizzled Class Receptor 4 |
| *FZD5* | Frizzled Class Receptor 5 |
| *FZD6* | Frizzled Class Receptor 6 |
| *FZD7* | Frizzled Class Receptor 7 |
| *FZD8* | Frizzled Class Receptor 8 |
| *FZD9* | Frizzled Class Receptor 9 |
| *FZD10* | Frizzled Class Receptor 10 |
| *GSK3B* | Glycogen Synthase Kinase 3 Beta |
| *LGR4* | Leucine Rich Repeat Containing G Protein-Coupled Receptor 4 |
| *LGR5* | Leucine Rich Repeat Containing G Protein-Coupled Receptor 5 |
| *LGR6* | Leucine Rich Repeat Containing G Protein-Coupled Receptor 6 |
| *LRP5* | Low-density lipoprotein receptor-related protein 5 |
| *LRP6* | Low-density lipoprotein receptor-related protein 6 |
| *RNF43* | Ring finger protein 43 |
| *ZNRF3* | Zinc and Ring Finger 3 |

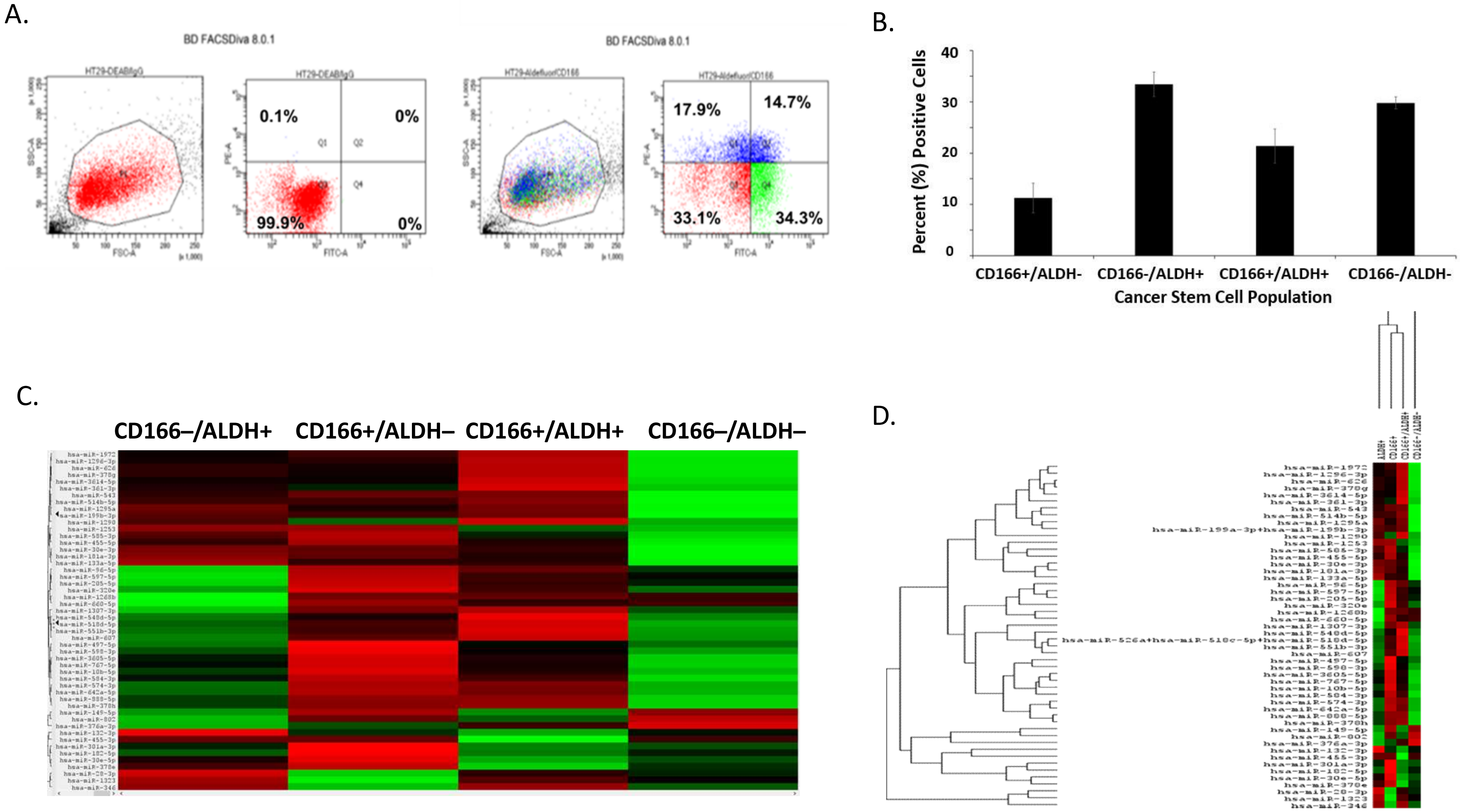
A.
BD FACSDiva 8.0.1
0.1%
0%
99.9%
0%
17.9%
14.7%
33.1%
34.3%
B.
Percent (%) Positive Cells
CD166+/ALDH-
CD166-/ALDH+
CD166+/ALDH+
CD166-/ALDH-
Cancer Stem Cell Population
C.
CD166–/ALDH+
CD166+/ALDH–
CD166+/ALDH+
CD166–/ALDH–
D.

Figure S1

**E.**

**CD166+/ALDH- Top miRNAs**

| Name | *p*-Value | Ratio |
|---|---|---|
| hsa-miR-185-5p | 0.005335 | 1.52 |
| hsa-miR-34a-5p | 0.079482 | 1.84 |
| hsa-miR-28-5p | 0.097872 | 1.80 |
| hsa-miR-135b-5p | 0.13191 | 1.42 |
| hsa-miR-183-5p | 0.148579 | 1.23 |
| hsa-miR-1246 | 0.185359 | 1.91 |
| hsa-miR-1323 | 0.341565 | 2.08 |
| hsa-miR-26a-5p | 0.377387 | 1.21 |
| hsa-miR-196a-5p | 0.413641 | 1.31 |
| hsa-miR-1299 | 0.42265 | 1.22 |

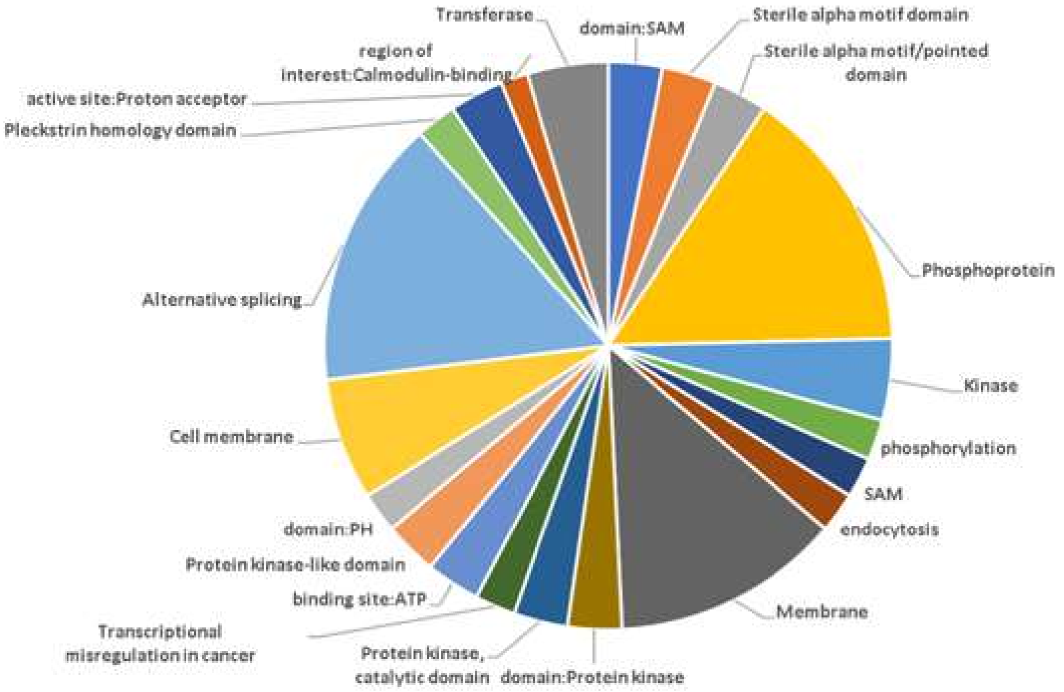

**F.**

**ALDH+/CD166- Top miRNAs**

| Name | *p*-Value | Ratio |
|---|---|---|
| hsa-miR-301a-3p | 0.024285 | 1.59 |
| hsa-miR-96-5p | 0.041435 | 1.61 |
| hsa-miR-29b-3p | 0.071412 | 1.18 |
| hsa-miR-429 | 0.084041 | 1.54 |
| hsa-miR-182-5p | 0.085607 | 1.54 |
| hsa-miR-660-5p | 0.100104 | 1.92 |
| hsa-miR-200a-3p | 0.100196 | 1.47 |
| hsa-miR-10a-5p | 0.104863 | 1.16 |
| hsa-miR-93-5p | 0.106804 | 1.25 |
| hsa-miR-194-5p | 0.107797 | 1.32 |

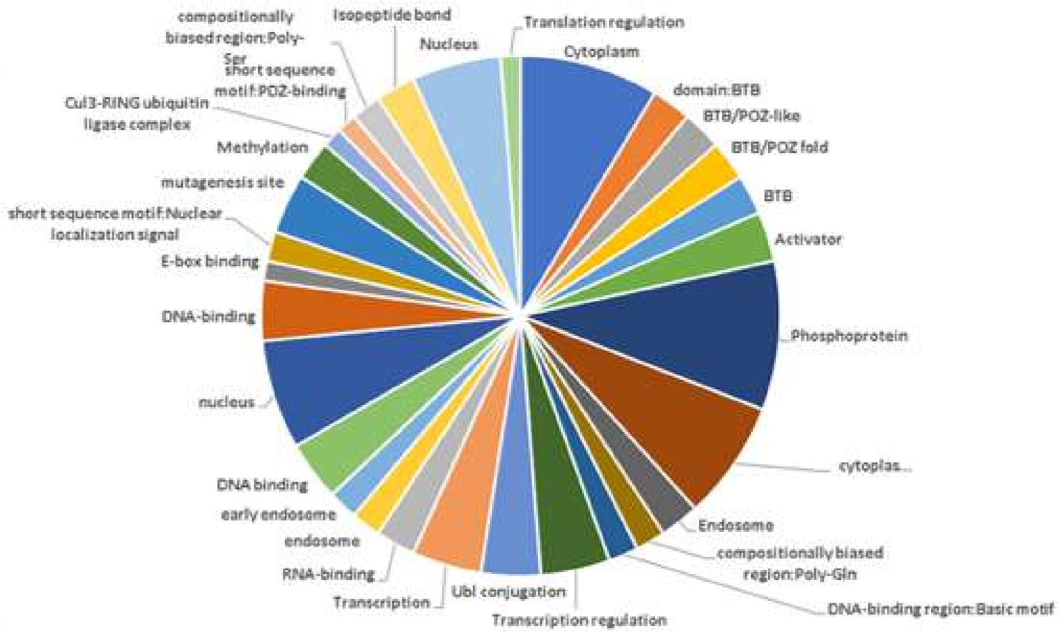

Figure S1

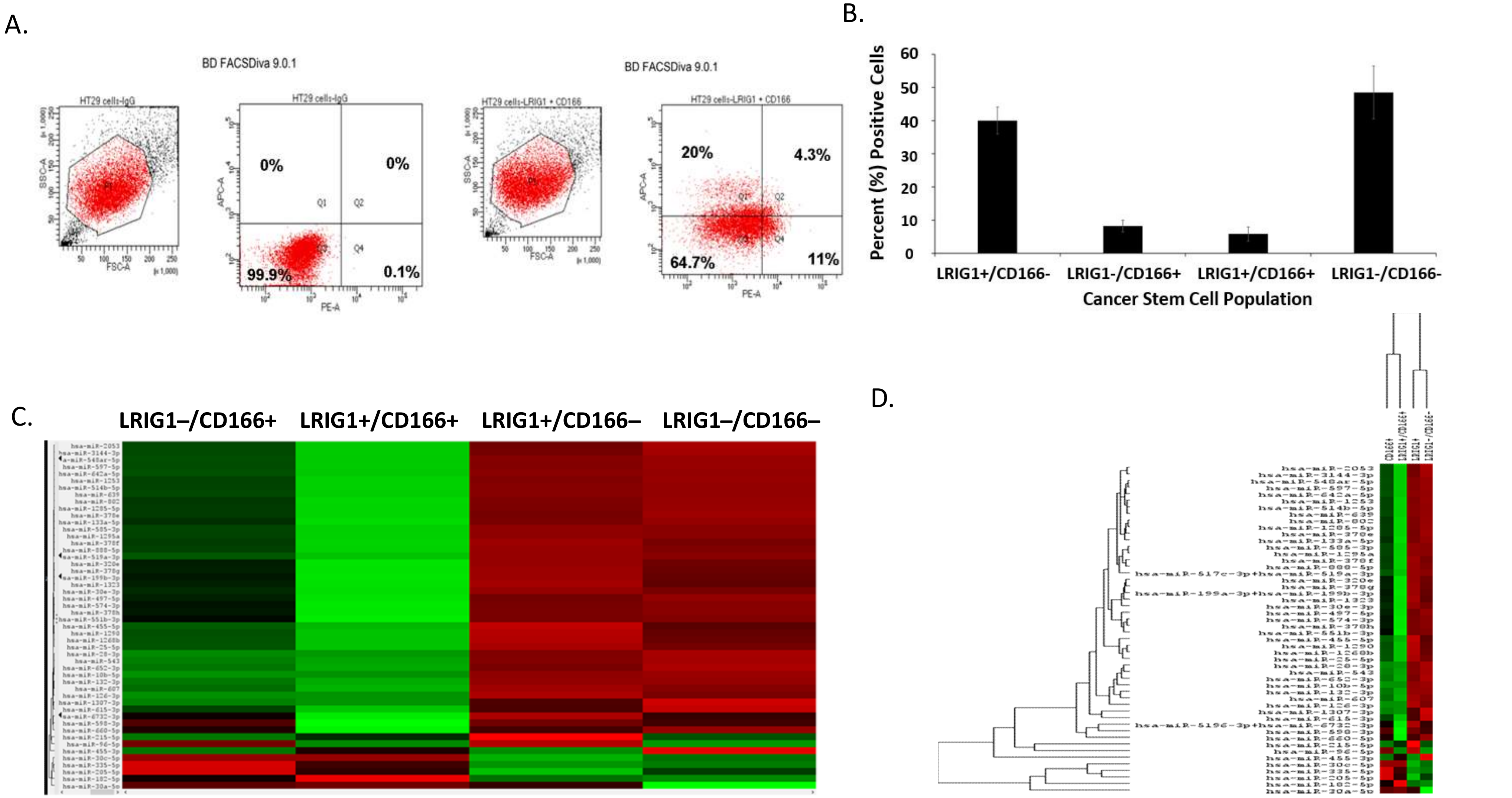
A.
BD FACSDiva 9.0.1
HT29 cells-IgG
HT29 cells-LRIG1 + CD166
0%
0%
99.9%
0.1%
20%
4.3%
64.7%
11%
B.
Percent (%) Positive Cells
LRIG1+/CD166-
LRIG1-/CD166+
LRIG1+/CD166+
LRIG1-/CD166-
Cancer Stem Cell Population
C.
LRIG1–/CD166+
LRIG1+/CD166+
LRIG1+/CD166–
LRIG1–/CD166–
D.

Figure S2

**E.**

**LRIG1+/CD166- Top miRNAs**

| Name | *p*-Value | Ratio |
|---|---|---|
| hsa-miR-374b-5p | 0.009738 | 1.39 |
| hsa-miR-205-5p | 0.010734 | 4.21 |
| hsa-let-7i-5p | 0.044163 | 1.58 |
| hsa-miR-192-5p | 0.04518 | 1.62 |
| hsa-miR-429 | 0.0745 | 1.64 |
| hsa-miR-106b-5p | 0.116775 | 1.43 |
| hsa-miR-590-5p | 0.126675 | 1.56 |
| hsa-miR-361-5p | 0.128048 | 1.18 |
| hsa-miR-27b-3p | 0.131583 | 1.4 |
| hsa-miR-374a-5p | 0.198531 | 1.58 |

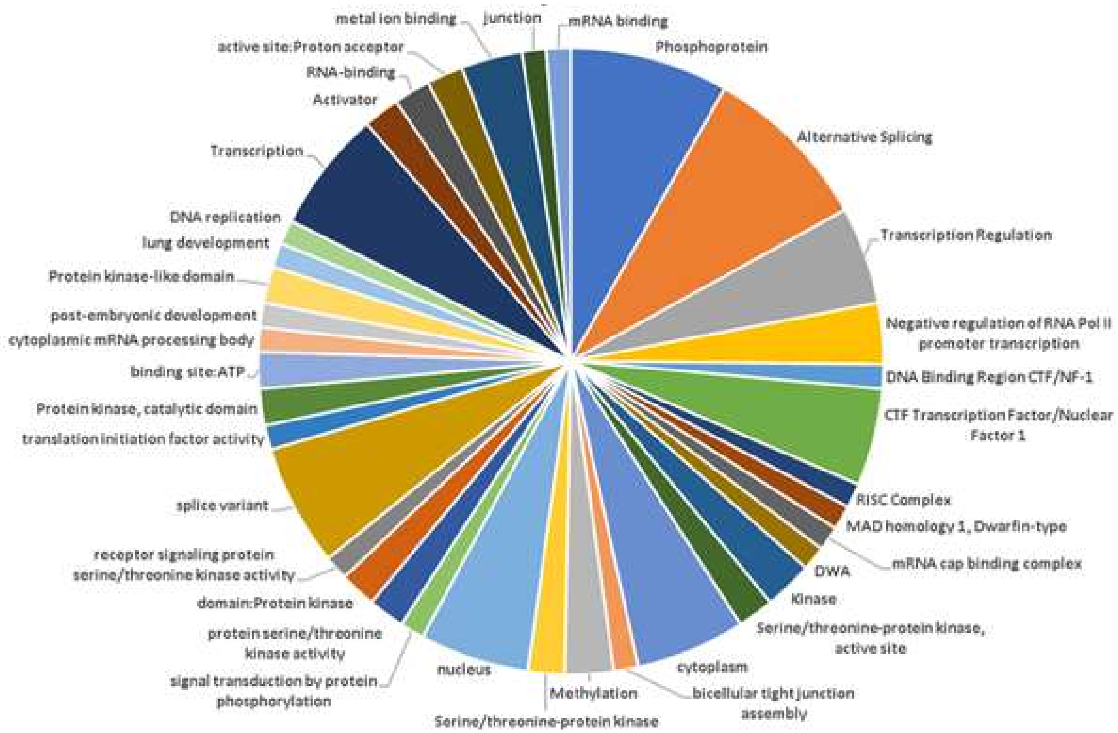

**F.**

**CD166+/LRIG1- Top miRNAs**

| Name | *p*-Value | Ratio |
|---|---|---|
| hsa-miR-608 | 0.004408 | 1.61 |
| hsa-miR-1200 | 0.006728 | 1.72 |
| hsa-miR-582-3p | 0.011376 | 2.00 |
| hsa-miR-548y | 0.012319 | 2.13 |
| hsa-miR-30a-3p | 0.014378 | 2.08 |
| hsa-miR-28-3p | 0.02381 | 1.64 |
| hsa-miR-517c-3p* | 0.024478 | 2.13 |
| hsa-miR-1302 | 0.025088 | 2.08 |
| hsa-miR-28-5p | 0.02628 | 2.04 |
| hsa-miR-628-3p | 0.027493 | 1.75 |

*hsa-miR-517c-3p+hsa-miR-519a-3p

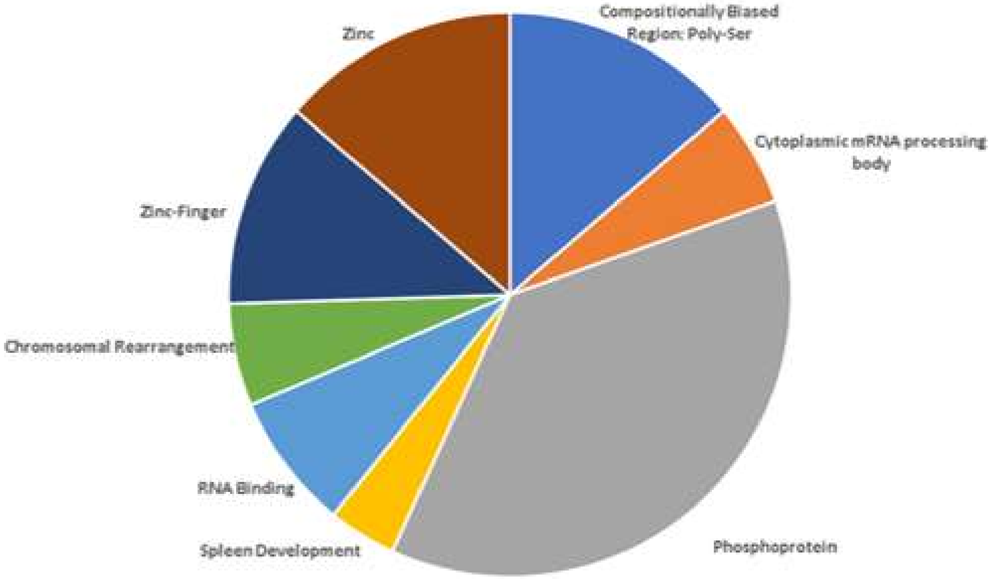

Figure S2

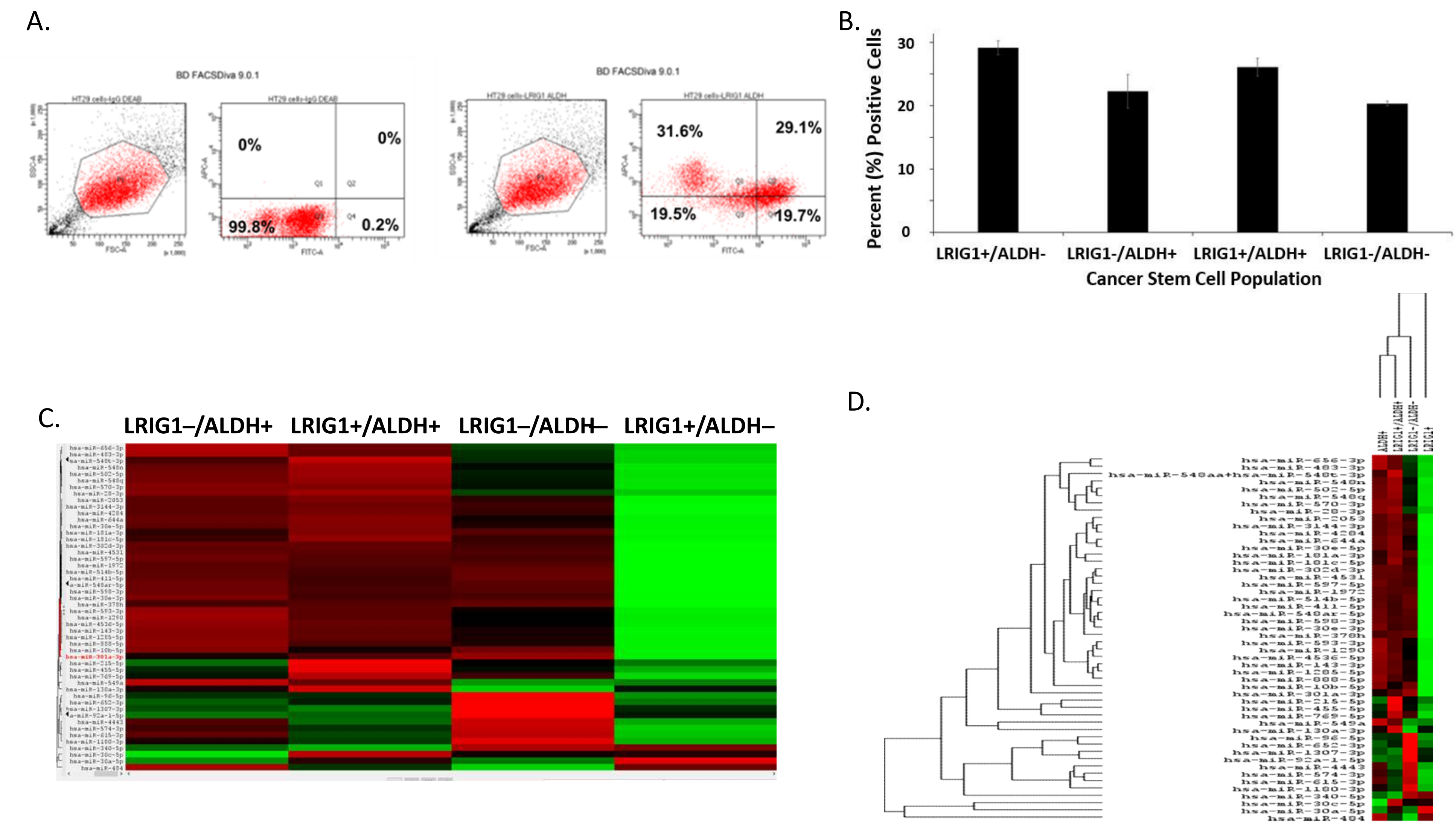
A.
BD FACSDiva 9.0.1
0%
0%
99.8%
0.2%
BD FACSDiva 9.0.1
31.6%
29.1%
19.5%
19.7%
B.
Percent (%) Positive Cells
30
20
10
0
LRIG1+/ALDH-
LRIG1-/ALDH+
LRIG1+/ALDH+
LRIG1-/ALDH-
Cancer Stem Cell Population
C.
LRIG1–/ALDH+
LRIG1+/ALDH+
LRIG1–/ALDH–
LRIG1+/ALDH–
D.

Figure S3

**E.** **LRIG1+/ALDH- Top miRNAs**

| Name | *p*-Value | Ratio |
|---|---|---|
| hsa-miR-5001-5p | 0.0029 | 2.41 |
| hsa-miR-193b-3p | 0.007301 | 2.03 |
| hsa-miR-1290 | 0.007807 | 3.22 |
| hsa-miR-208a-3p | 0.008697 | 3.06 |
| hsa-miR-510-3p | 0.00965 | 2.47 |
| hsa-miR-365b-5p | 0.010913 | 2.37 |
| hsa-miR-663a | 0.01348 | 3.70 |
| hsa-miR-1297 | 0.014174 | 2.19 |
| hsa-miR-205-5p | 0.014201 | 4.33 |
| hsa-miR-3614-5p | 0.015049 | 3.18 |

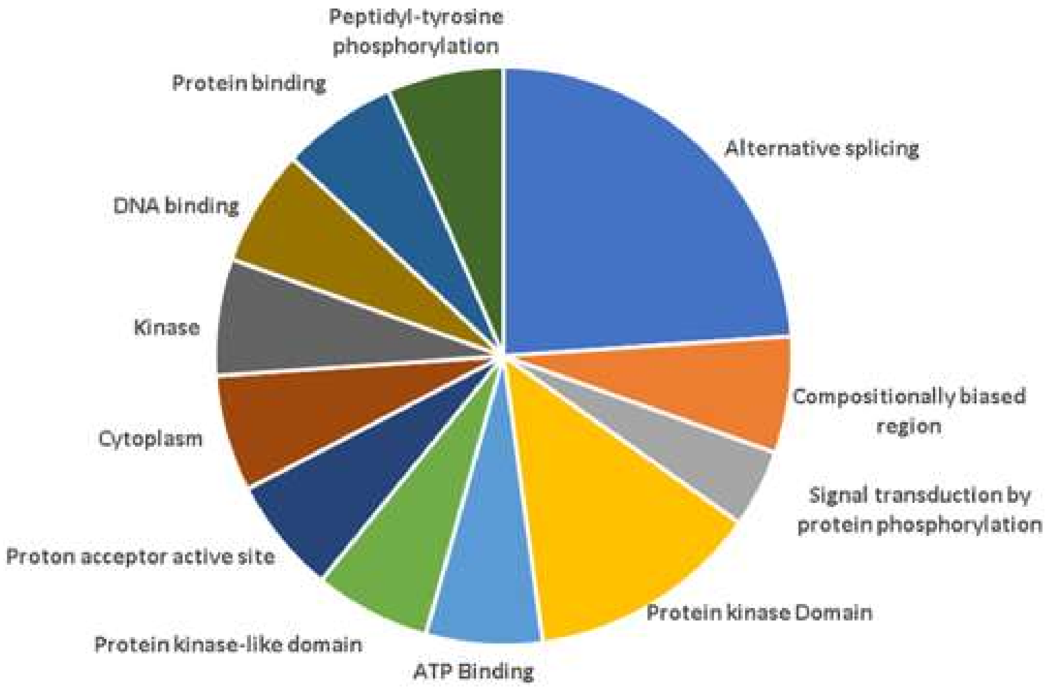

**F.** **ALDH+/LRIG1- Top miRNAs**

| Name | *p*-Value | Ratio |
|---|---|---|
| hsa-miR-148a-3p | 0.007872 | 1.64 |
| hsa-miR-3928-3p | 0.009459 | 1.12 |
| hsa-miR-194-5p | 0.012492 | 1.89 |
| hsa-miR-375 | 0.021317 | 1.61 |
| hsa-miR-22-3p | 0.056906 | 1.69 |
| hsa-miR-192-5p | 0.062684 | 1.56 |
| hsa-miR-185-5p | 0.064618 | 1.32 |
| hsa-miR-19a-3p | 0.068956 | 1.96 |
| hsa-miR-140-5p | 0.071816 | 1.75 |
| hsa-miR-27b-3p | 0.078661 | 1.37 |

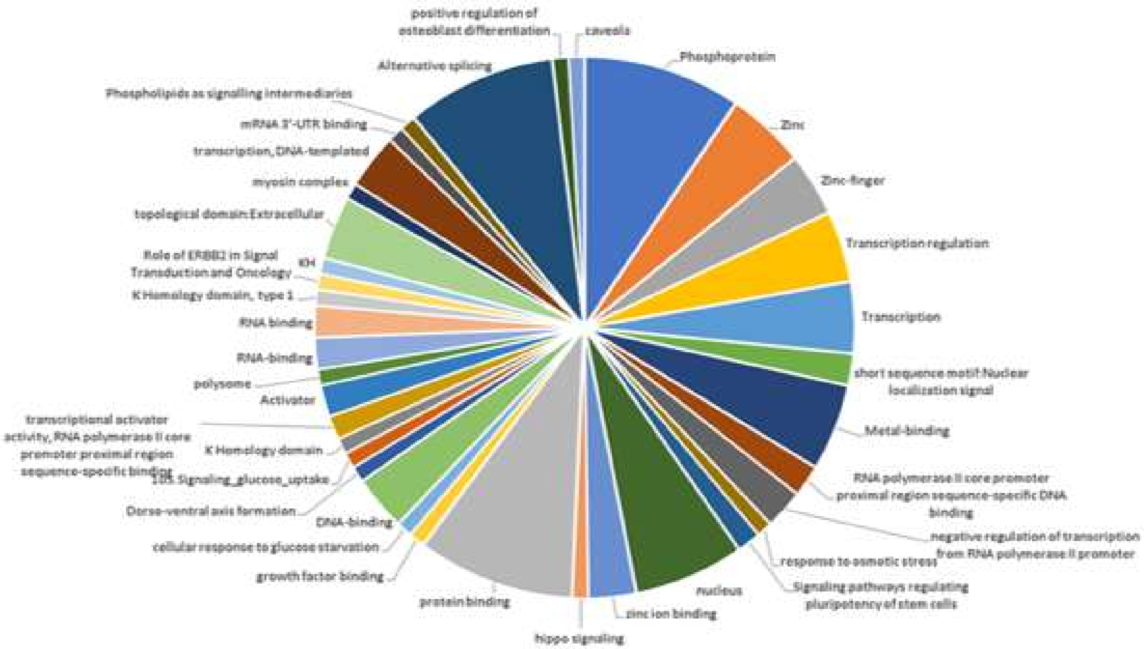

Figure S3

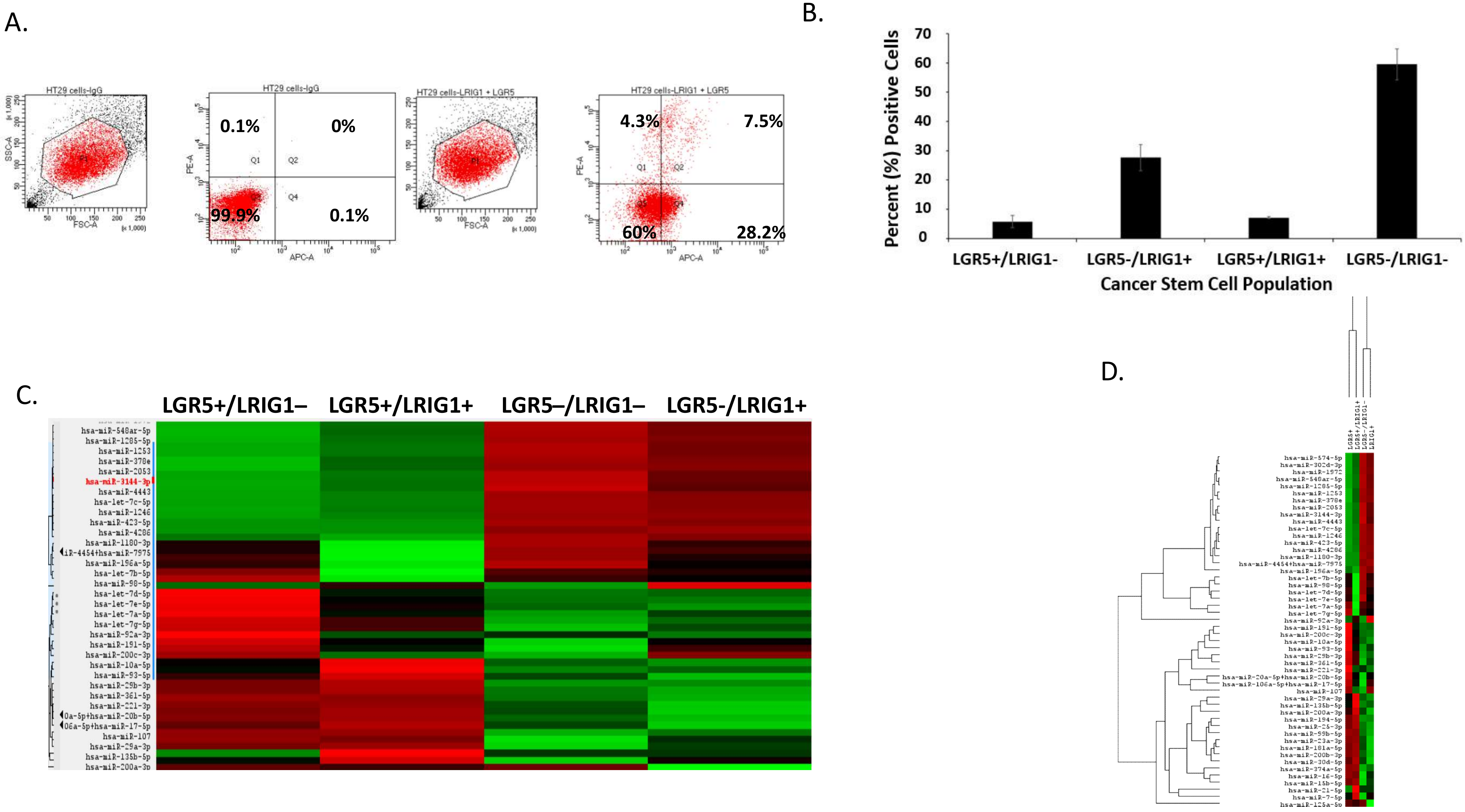
A.
HT29 cells-IgG
SSC-A
FSC-A
HT29 cells-IgG
0.1%
0%
99.9%
0.1%
PE-A
APC-A
HT29 cells-LRIG1 + LGR5
HT29 cells-LRIG1 + LGR5
4.3%
7.5%
60%
28.2%
B.
Percent (%) Positive Cells
70
60
50
40
30
20
10
0
LGR5+/LRIG1-
LGR5-/LRIG1+
LGR5+/LRIG1+
LGR5-/LRIG1-
Cancer Stem Cell Population
C.
LGR5+/LRIG1–
LGR5+/LRIG1+
LGR5–/LRIG1–
LGR5-/LRIG1+
D.

Figure S4

**E.**

**LGR5+/LRIG1- Top miRNAs**

| Name | *p*-Value | Ratio |
|---|---|---|
| hsa-miR-455-3p | 0.000516 | 5.626198 |
| hsa-miR-424-5p | 0.001786 | 7.473064 |
| hsa-miR-454-3p | 0.00185 | 3.375876 |
| hsa-miR-544a | 0.001903 | 9.050518 |
| hsa-miR-935 | 0.001946 | 9.763292 |
| hsa-miR-874-5p | 0.002336 | 11.21283 |
| hsa-miR-219a-2-3p | 0.002448 | 13.2283 |
| hsa-miR-218-5p | 0.002587 | 9.866657 |
| hsa-miR-532-5p | 0.002709 | 6.301194 |
| hsa-miR-1972 | 0.002842 | 24.14922 |

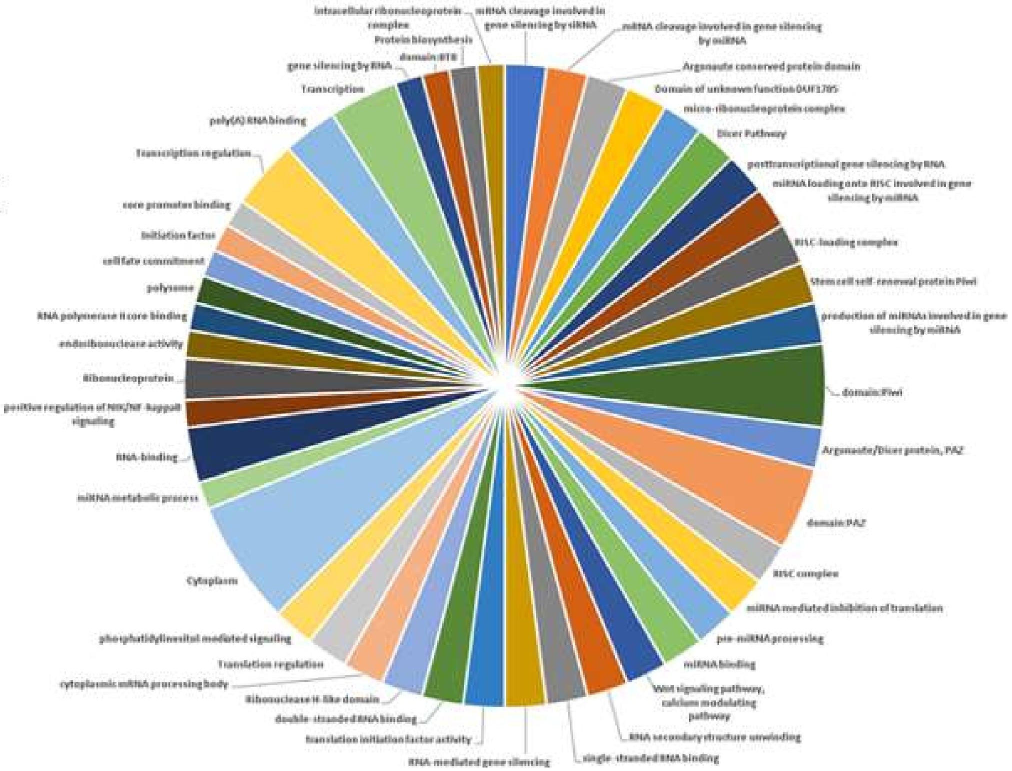

**F.**

**LRIG1+/LGR5- Top miRNAs**

| Name | *p*-Value | Ratio |
|---|---|---|
| hsa-miR-200b-3p | 0.006559 | 2.129456 |
| hsa-miR-194-5p | 0.010642 | 2.207357 |
| hsa-miR-23a-3p | 0.032224 | 1.784871 |
| hsa-miR-29a-3p | 0.034529 | 1.311528 |
| hsa-miR-10a-5p | 0.06363 | 2.184755 |
| hsa-miR-30d-5p | 0.071142 | 1.691372 |
| hsa-miR-15b-5p | 0.092976 | 1.587857 |
| hsa-miR-374a-5p | 0.107158 | 2.051671 |
| hsa-miR-191-5p | 0.120353 | 1.675972 |
| hsa-miR-361-5p | 0.128485 | 1.191644 |

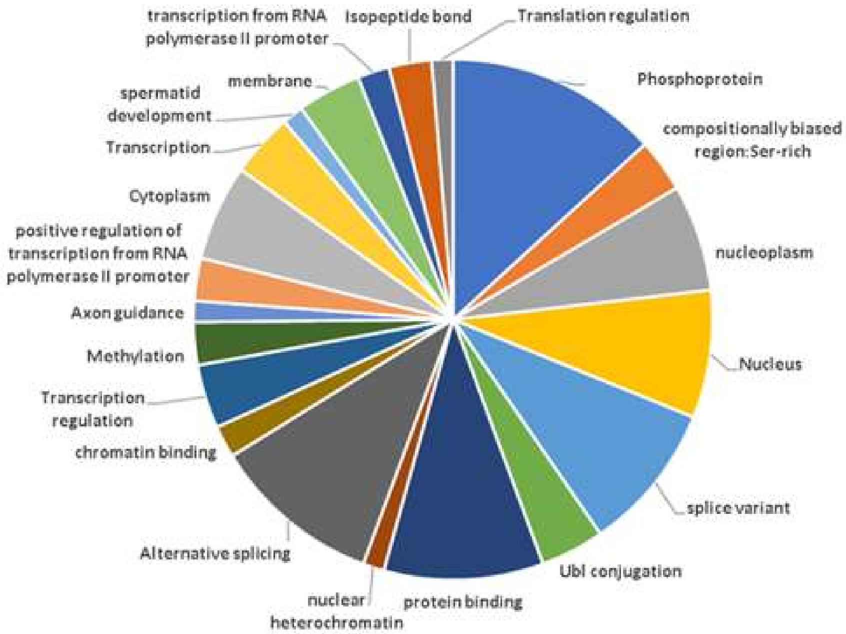

Figure S4

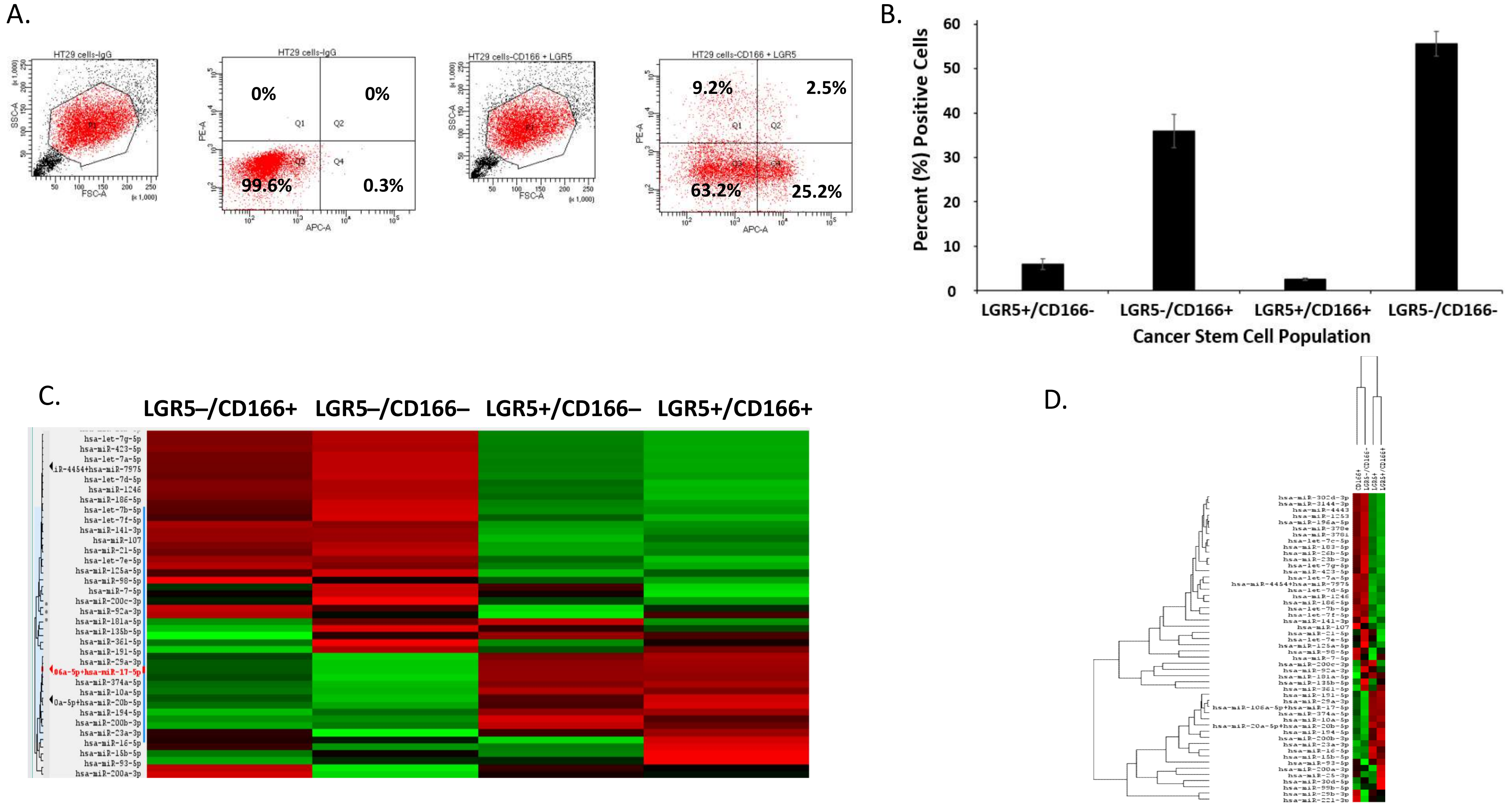
A.
HT29 cells-IgG
HT29 cells-CD166 + LGR5
0%
0%
99.6%
0.3%
9.2%
2.5%
63.2%
25.2%
B.
Percent (%) Positive Cells
LGR5+/CD166-
LGR5-/CD166+
LGR5+/CD166+
LGR5-/CD166-
Cancer Stem Cell Population
C.
LGR5–/CD166+
LGR5–/CD166–
LGR5+/CD166–
LGR5+/CD166+
D.

Figure S5

## E.

#### LGR5+/CD166- Top miRNAs

| Name | *p*-Value | Ratio |
|---|---|---|
| hsa-miR-4454* | 0.002637 | 8.333 |
| hsa-miR-664b-5p | 0.005766 | 6.667 |
| hsa-miR-196a-5p | 0.005853 | 1.370 |
| hsa-miR-3918 | 0.010346 | 5.556 |
| hsa-miR-1204 | 0.013975 | 10.000 |
| hsa-miR-376a-2-5p | 0.014634 | 6.250 |
| hsa-miR-362-3p | 0.01532 | 9.091 |
| hsa-miR-142-5p | 0.015663 | 5.556 |
| hsa-miR-1908-5p | 0.017958 | 10.000 |
| hsa-miR-142-3p | 0.019367 | 3.226 |

*hsa-miR-4454+hsa-miR-7975

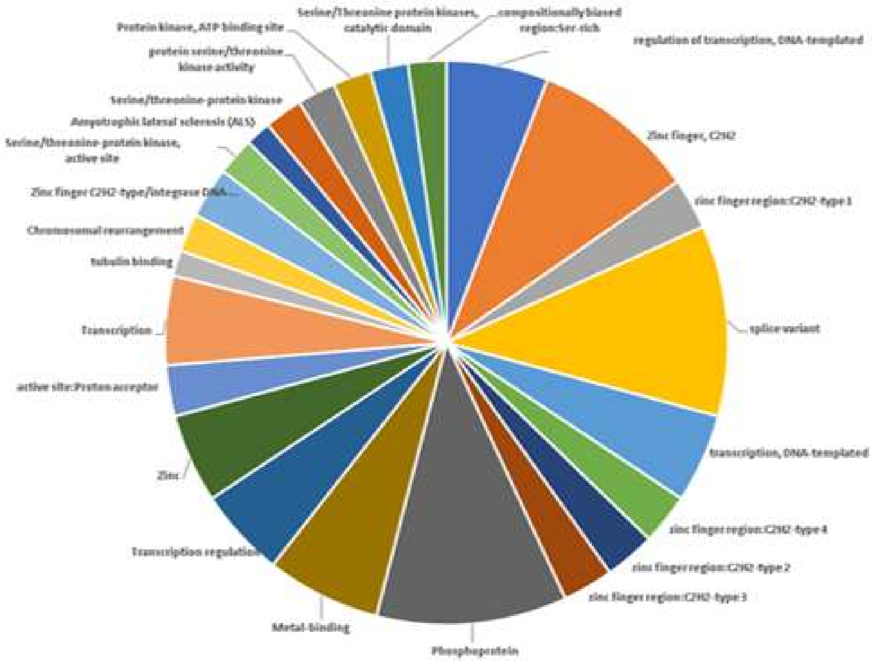

## F.

#### CD166+/LGR5- Top miRNAs

| Name | *p*-Value | Ratio |
|---|---|---|
| hsa-miR-15b-5p | 0.034841 | 2.06 |
| hsa-miR-16-5p | 0.055473 | 2.04 |
| hsa-miR-29a-3p | 0.143658 | 1.41 |
| hsa-miR-194-5p | 0.165687 | 1.44 |
| hsa-miR-20a-5p* | 0.174565 | 1.48 |
| hsa-miR-1283 | 0.212371 | 3.81 |
| hsa-miR-181a-5p | 0.213746 | 1.46 |
| hsa-miR-23a-3p | 0.25129 | 1.37 |
| hsa-miR-106a-5p** | 0.26257 | 1.17 |
| hsa-let-7i-5p | 0.267539 | 11.00 |

*hsa-miR-20a-5p+hsa-miR-20b-5p, **hsa-miR-106a-5p+hsa-miR-17-5p

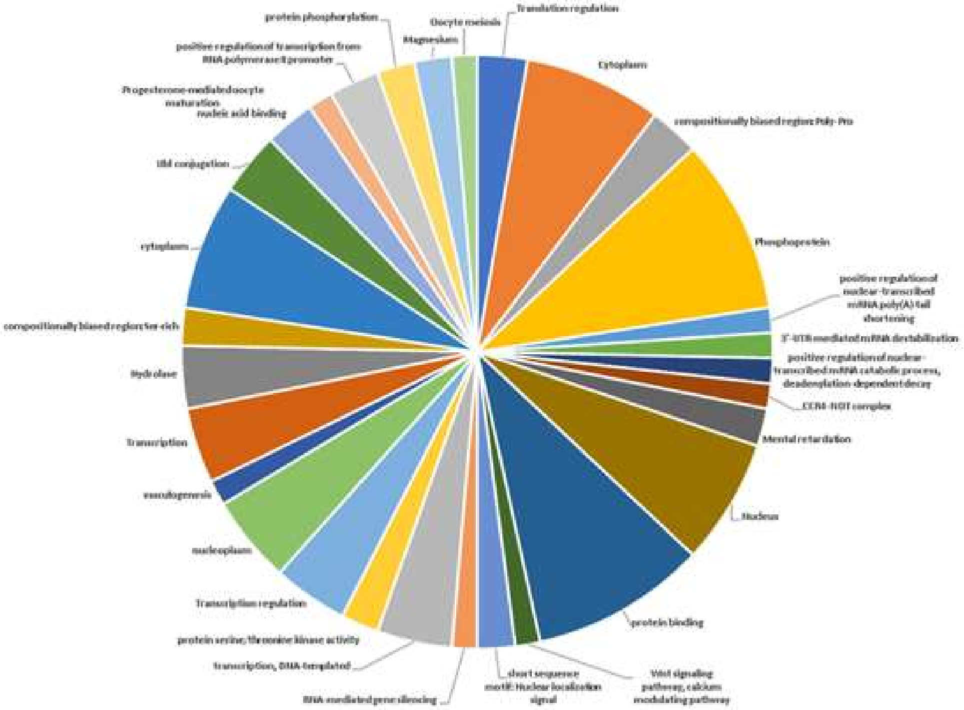

Figure S5
